# PP1 PNUTS binds the “restrictor” and dephosphorylates RNA pol II CTD Ser5 to stimulate transcription termination

**DOI:** 10.1101/2024.07.12.603302

**Authors:** Benjamin Erickson, Roman Fedoryshchak, Nova Fong, Ryan Sheridan, Keira Y. Larson, Anthony J. Saviola, Stephane Mouilleron, Kirk C. Hansen, Richard Treisman, David L. Bentley

## Abstract

The restrictor, ZC3H4/WDR82, is the major termination factor for antisense transcription from bidirectional promoters, but its mechanism is poorly understood. We report that ZC3H4/WDR82 co-purifies with PP1 phosphatase and PP1 phosphatase nuclear targeting subunit, PNUTS, which binds directly to the WDR82 subunit of restrictor. AlphaFold predicts a quaternary complex, PPWZ, in which <u>P</u>P1-associated <u>P</u>NUTS and <u>Z</u>C3H4 both contact <u>W</u>DR82. To investigate the role of protein dephosphorylation in PPWZ activity, we expressed a substrate trap comprising inactive PP1^H66K^ linked to the PNUTS C-terminus. PP1^H66K^-PNUTS binds pol II large subunit and nuclear exosome components. PP1^H66K^-PNUTS, but not PP1^WT^-PNUTS, functions as a dominant-negative inhibitor of antisense termination and CTD Ser5 dephosphorylation. Both these activities require the PNUTS WDR82 binding domain that interacts with restrictor. We show that CTD Ser5 hyperphosphorylation is associated with higher processivity and reduced pausing that would counteract termination, and propose that Ser5 dephosphorylation by PPWZ is coupled to termination. In summary, we identify the PP1 phosphatase activity of the PPWZ complex as essential for terminator function and propose that this heterotetramer is the physiologically relevant form of restrictor.

## Introduction

Most promoters in multicellular organisms are to some extent bidirectional and give rise to both sense and divergent antisense transcription which must eventually terminate to prevent interference with neighboring genes and collisions with replication forks. Sense transcription of most protein coding genes terminates via the “torpedo” mechanism that couples termination to RNA cleavage at the polyA site. This permits loading of the 5’3’ RNA exonuclease “torpedo” Xrn2/Rat1 which facilitates polymerase release (Connelly and Manley 1988; Kim et al. 2004; West et al. 2004; Zeng et al. 2024). In contrast, antisense transcription from promoters and bidirectional transcription from enhancers generate unstable non-coding RNAs that are largely terminated by mechanisms independent of cleavage/polyadenylation (CPA) (Andersson et al. 2015; Austenaa et al. 2015). An important type of polyA site independent termination depends on the “restrictor” a heterodimer of the WD40 domain protein WDR82 and the Zn finger protein ZC3H4 that is conserved between Drosophila and humans (Austenaa et al. 2015; Brewer-Jensen et al. 2015). WDR82-ZC3H4 interacts with mRNA 5’ Cap Binding Complex-associated factor ARS2/SRRT (Estell et al. 2023; Rouvière et al. 2023) that also binds the nuclear exosome implicated in coupled termination and RNA degradation (Tudek et al. 2014; Iasillo et al. 2017). How restrictor activity is directed to antisense transcripts remains unclear; it has been suggested that binding to sense transcripts is antagonized by SET1 histone methyltransferases, which bind WDR82 mutually exclusively with ZC3H4 (Estell et al. 2023; Hughes et al. 2023).

RNA pol II CTD phosphorylation and dephosphorylation control transitions within the transcription cycle (Buratowski 2009), but how they contribute specifically to elongation-termination transitions is not well understood. The Ser5 residues of the pol II CTD heptad repeats (YSPTSPS) are phosphorylated by TFIIH-associated CDK7 and dephosphorylated by PP1/PNUTS complexed with WDR82 and Tox4 (Lee and Skalnik 2008; Lee et al. 2010; Ciurciu et al. 2013; Landsverk et al. 2020) and Integrator/PP2A(Huang et al. 2020; Vervoort et al. 2021; Hu et al. 2023). At 5’ ends, Ser5 dephosphorylation by Integrator/PP2A is associated with attenuation of sense transcripts (Huang et al. 2020; Vervoort et al. 2021; Hu et al. 2023). At 3’ ends, Ser5 dephosphorylation precedes polyA site-dependent transcription termination. PP1, PNUTS, WDR82 and ZC3H4 all bind to the Ser5 phosphorylated CTD (Ebmeier et al. 2017; Wu et al. 2018), and consistent with a role in co-transcriptional dephosphorylation, PP1 and PNUTS localize on transcribed genes (Verheyen et al. 2015; Cortazar et al. 2019; Cossa et al. 2020). How CTD Ser5 dephosphorylation might facilitate termination is unclear.

The PP1 phosphatase nuclear targeting subunit, PNUTS, functions in both polyA site-independent and -dependent termination. PNUTS co-purifies with mammalian cleavage polyadenylation complexes (Shi et al. 2009), and its knockdown inhibits termination in several contexts (Kieft et al. 2020; Devlin et al. 2022) including divergent antisense transcription from bidirectional promoters (Austenaa et al. 2015; Estell et al. 2023). PNUTS has short linear motifs (SLIMs) that bind to PP1(Choy et al. 2014) as well as a TFIIS homology domain, an RNA binding RGG domain (Kreivi et al. 1997; Kim et al. 2003) and a WDR82-interaction domain that is conserved from yeast to humans (Lee et al. 2010; Vanoosthuyse et al. 2014; Benjamin et al. 2021). PNUTS positively affects PP1 activity by facilitating substrate recruitment but also constrains its specificity by occupying one of the three substrate binding grooves on the PP1 surface (Choy et al. 2014). Hence interpretation of PNUTS knockdown phenotypes is not straightforward. Exactly how PNUTS functions in termination by the restrictor, and whether PP1 dependent dephosphorylation has a role in this process are unclear. It has been suggested that it plays an indirect role, possibly by slowing pol II elongation (Estell et al. 2023; Russo et al. 2023) as it does at polyA sites (Parua et al. 2018; Cortazar et al. 2019). Previous studies showed that ZC3H4 co-immunoprecipitated with WDR82 but not with PNUTS, leading to the proposal that WDR82 interaction with PNUTS and ZC3H4 is mutually exclusive (Austenaa et al. 2021), although PNUTS/ZC3H4 interaction can be detected in vivo by proximity labelling (Estell et al. 2023; Russo et al. 2023). PNUTS and exosome knock-down mimic the effects of restrictor inactivation, but no specific molecular interaction or enzymatic activity of any interacting factor is known to be required for termination by restrictor.

The subcellular targeting and substrate specificity of PP1 holoenzymes is determined by the association of the PP1 catalytic subunit with different PP1 interacting proteins (PIPs) (Heroes et al. 2013), which has hampered the development of holoenzyme-specific small molecule modulators (Cossa et al. 2021). This difficulty can be addressed by the use of chimeric fusions. We previously showed that a fusion of PP1 with the Phactr/PIP retains the specificity of the PP1/Phactr holoenzyme (Fedoryshchak et al. 2020). Here we use a similar approach to assess the function of PP1/PNUTS. Expression of a PP1^WT^-PNUTS fusion has only a modest effect on transcription but remarkably inactive PP1^H66K^-PNUTS (Zhang et al. 1996) strongly inhibited termination of divergent antisense pol II transcription and dephosphorylation of pol II resulting in Ser5 hyperphosphorylation. Moreover PP1^H66K^-PNUTS interacted directly with the restrictor through the WDR82 binding domain (WBD) of PNUTS, and this interaction was required for its dominant negative effect on CTD dephosphorylation and termination. Together these results suggest that the phosphatase activity of PP1/PNUTS bound to restrictor couples termination to dephosphorylation of pol II CTD Ser5.

## Results

### PNUTS binds to the restrictor

We investigated factors interacting with PNUTS by Mass Spec identification of proteins immunoprecipitated with epitope tagged PNUTS from benzonase treated nuclear extracts (Figure 1A, B). Avitag-PNUTS was over-expressed in HEK293 cells from a doxcycyline inducible lentiviral vector. As expected, PNUTS IP recovered PP1 catalytic (PPP1CA) and regulatory subunits including the inhibitor PPP1R8, PPP1R2 and PPP1R9B (spinophilin) (Heroes et al. 2013) as well as Tox4 (Lee et al. 2010) (Figure1C). Pol II large subunit was also modestly enriched in the PNUTS IP consistent with previous results (Ebmeier et al. 2017; Wu et al. 2018). Remarkably, both the WDR82 and ZC3H4 subunits of restrictor were among the most highly enriched proteins co-purifying with PNUTS (Figure 1C). This observation is surprising as ZC3H4 and PNUTS were previously thought to bind WDR82 in a mutually exclusive manner (Austenaa et al. 2021).

**Figure 1.**
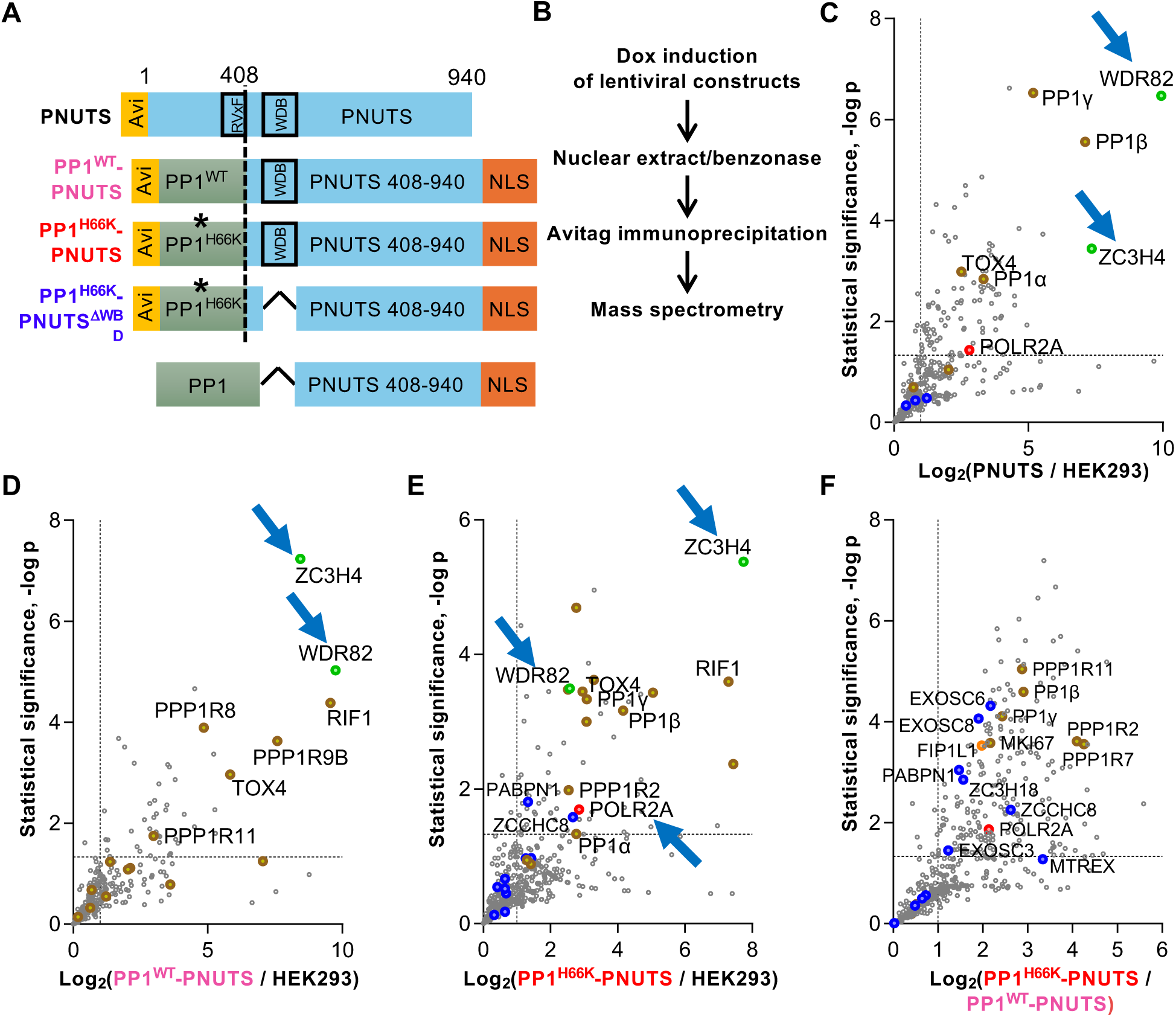
PNUTS binds restrictor via its WDR82 binding domain. **A.** Diagrams of Avitag constructs inducibly expressed from lentiviral vectors in HEK293 used for IP-MS. **B.** IP-MS strategy. **C.** PNUTS co-IPs with both subunits of restrictor (blue arrows). Full length Avitag-PNUTS 1-940 was immunoprecipitated and Log2 Fold Change relative to anti-Avitag IP’s from parental HEK293 cells plotted. 4-6 replicates per condition. **D.** PP1^WT^-PNUTS co-immunoprecipitates with both subunits of restrictor. **E.** PP1^H66K^-PNUTS co-immunoprecipitates with both subunits of restrictor and pol II large subunit (blue arrows). **F.** Pol 2 large subunit and nuclear exosome associated factors (blue labels) are specifically enriched in IP’s of PP1^H66K^-PNUTS relative to PP1^WT^-PNUTS.

### Candidate substrates for PP1-PNUTS are linked to restrictor termination

We and others have used chimeric fusions of PP1-with its co-factors to accurately reproduce the substrate specificity of PP1 holoenzymes and to trap substrates (Wu et al. 2018; Fedoryshchak et al. 2020). We constructed a fusion between Avitag PP1α (1-304) and the C-terminus of PNUTS (408-940) (Figure 1A) that was inducibly expressed in HEK293 cells from a lentiviral vector. In this chimera the conserved PP1 interaction motif RVxF (398-401) of PNUTS is deleted and the two proteins are connected by a flexible linker (SGSGS). IP-MS analysis of the PP1^WT^-PNUTS fusion revealed that, similar to full-length PNUTS, two of the most significantly enriched co-purifying proteins are the WDR82 and ZC3H4 subunits of restrictor along with known PP1 and PNUTS interactors PPP1R8, PPP1R9B, Tox4, and RIF1 (Figure 1D).

To identify candidate substrates of PP1/PNUTS we designed a catalytically inactive PP1^H66K^-PNUTS fusion in which a key metal binding residue is substituted. We compared the interactomes of PP1^WT^- and PP1^H66K^-PNUTS fusion proteins seeking proteins that were recovered more effectively by the catalytically inactive mutant (Wu et al. 2018). IP-MS revealed that PP1^H66K^-PNUTS, like full-length PNUTS and the PP1^WT^-PNUTS fusion, also effectively pulled down both restrictor subunits along with PP1 regulatory subunits and PNUTS interactors PPP1R2 (Inhibitor-2), PPP1R8, PPP1R7, PPP1R11, PPP1R9A/B, Tox4, MK167 and RIF1 (Figure 1E). In agreement with previous work (Wu et al. 2018), the pol II large subunit POLR2A, whose CTD Ser5 residues are known substrates of PP1/PNUTS, is significantly enriched for binding to PP1^H66K^-PNUTS relative to PP1^WT^-PNUTS (Figure 1F). Notable other candidate substrates specifically enriched for binding to PP1^H66K^-PNUTS included the putative WDR82-associated termination factor FIP1L1 (Spencley et al. 2023) and subunits of nuclear exosome complexes that are implicated in restrictor function (Estell et al. 2023; Rouvière et al. 2023). These include the exonuclease MTREX (Mtr4), the ARS2 interacting phospho-protein ZC3H18(Winczura et al. 2018), the NEXT subunit ZCCHC8, the PAXT-associated PABPN1 and other polyA binding proteins, as well as the core exosome subunits EXOSC3, 6 and 8 (Figure 1F, Supp. Table S1). Together these results confirm the pol II large subunit as a PP1/PNUTS substrate and furthermore suggest several nuclear exosome associated proteins as additional potential targets.

### PNUTS binds restrictor via its WDR82 binding domain

The strong co-immunoprecipitation of PNUTS and PP1-PNUTS fusions with restrictor was unexpected as previous work suggested a loose interaction detected only by long-range proximity labelling (Austenaa et al. 2021; Estell et al. 2023; Russo et al. 2023). We asked whether restrictor contact with PNUTS is direct by determining whether it required the WDR82 binding domain (WBD). IP MS interactome analysis was performed on a fusion protein PP1^H66K^-PNUTS^ΔWBD^ comprising PP1(H66K) with a fragment of PNUTS (408-940) +NLS deleted for the WBD (residues 455-523) (Fig. 1A) that was expressed in HEK293 cells. Comparison of the interactomes showed that, as expected, WDR82 binding was significantly greater for the H66K-PNUTS control than the H66K-PNUTS^ΔWDR^ mutant. Importantly, ZC3H4 binding was also strongly enhanced in the H66K-PNUTS relative to the H66K-PNUTS^ΔWDR^ mutant (Fig. 2A) showing that ZC3H4 binding to PNUTS is mediated by its interaction with WDR82. This result was confirmed by Western blotting of immunoprecipitated PP1^H66K^-PNUTS and PP1^H66K^-PNUTS^ΔWBD^ fusion proteins (Figure 2B). Together these results establish that a PP1-PNUTS fusion binds to restrictor via its WBD and that this interaction is therefore almost certainly direct. In addition we found that association of PP1^H66K^-PNUTS with nuclear exosome associated proteins EXOSC3, 6, 8, and ZC3H18 was also dependent on WDR82 binding (Figure 2A). These results are therefore consistent with interaction between PNUTS-bound restrictor and the nuclear exosome (Estell et al. 2023; Rouvière et al. 2023).

**Figure 2.**
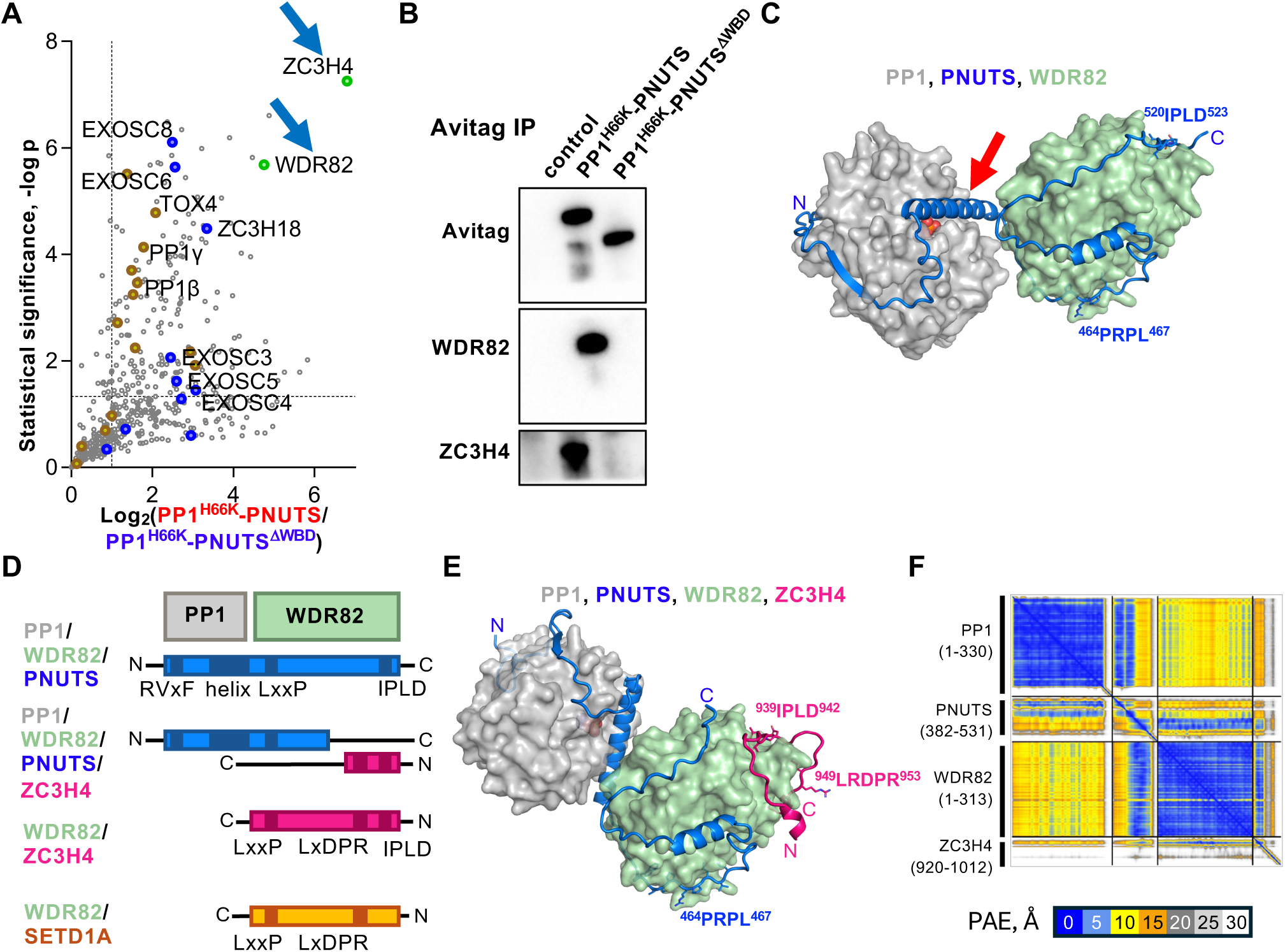
WDR82 contacts PNUTS and ZC3H4 in a predicted PPWZ complex of PP1/PNUTS/WDR82/ZC3H4. **A.** The ZC3H4 and WDR82 subunits of restrictor are specifically enriched in IP’s of Avitag-PP1^H66K^-PNUTS relative to Avitag-PP1^H66K^-PNUTS^△WBD^. IP-MS as in Figure1. **B.** IP Western of Avitag PP1^H66K^-PNUTS and PP1^H66K^-PNUTS^△WBD^. Note restrictor binding is dependent on the PNUTS WBD. **C.** AlphaFold prediction of a trimeric complex of PP1/PNUTS with WDR82. In 19/25 models including this one (ranked 3, see Figure S1A for the predicted aligned error plot) a helix of PNUTS (red arrow) occupies the active site of PP1. In 6/25 models this helix is not engaged with PP1 (Figure S1B). The IPLD and LxxP motifs in PNUTS predicted to contact WDR82 are indicated. **D.** Summary of Alphafold predicted SLIM’s (dark boxes) in PNUTS, ZC3H4 and SET1 that contact PP1 or WDR82 in the complexes listed at left. **E, F.** AlphaFold prediction and the predicted aligned error plot of the PPWZ complex of PP1/PNUTS/WDR82/ZC3H4. Sequences of SLIMs in ZC3H4 and PNUTS that mediate compatible interactions with WDR82 are marked in pink and blue. Note the IPLD motif in PNUTS is displaced from the surface of WDR82 by a competing motif in ZC3H4.

### PPWZ, a predicted complex of PP1/PNUTS/WDR82/ZC3H4

To gain insight into how PP1/PNUTS might interact with restrictor we performed AlphaFold2 modelling. Models of the PP1/PNUTS/WDR82 complex yielded high confidence predictions of PNUTS residues 397-451 binding to PP1 and residues 453-524 binding to WDR82 as indicated by the predicted aligned error (PAE) plot (Figures 2C, S1A). The PP1/PNUTS/WDR82 models are of two classes that differ in the interaction between PP1 and PNUTS. In 19 of 25 models, PNUTS residues 429-452 form an apparently auto-inhibitory helix that occludes the active site of PP1 (Figure 2C) and closely mimics the structure of PP1 inhibitor 2 complexed with PP1 (Hurley et al. 2007). In the other 6 output models, which closely conform with the structure of PP1 complexed with a fragment of PNUTS (394-433)(Choy et al. 2014), this helix is free (Figure S1B). The functional significance of these two alternative modes of PP1-PNUTS interaction for regulation of phosphatase activity remains to be investigated. In both types of model, PNUTS and WDR82 are predicted to interact through multiple close interactions along a trajectory of 80 residues which zig-zags across the WDR82 surface. These contacts include those made by PNUTS sequences ^461^WVCPRPLVL^469^ and ^520^IPLD^523^ that form LxxP and IPLD motifs discussed below (Figure 2C, D, S1E, F).

We next used Alphafold to predict the structure of a putative tetrameric PP1/PNUTS/WDR82/ZC3H4 “PPWZ” complex. In this predicted complex the PNUTS ^520^IPLD^523^-WDR82 interaction is replaced by the analogous ^939^IPLD^942^ motif from Z3CH4 (Figure S1E). ZC3H4 is also predicted to interact with WDR82 through its ^949^LRDPR^953^ sequence (Figure 2D-F) as shown previously ^8^. These interactions are also predicted by Alphafold for the complex of WDR82/ZC3H4 without PP1/PNUTS (Figure S1C, D), which includes additional interactions mediated by the ZC3H4 ^63^ELEDD^67^ and ^966^VPLSKPSF^971^ sequences (Figure 2D, S1F). The ZC3H4 ^63^ELEDD^67^ motif was previously shown to also bind ARS2 (Rouvière et al. 2023). Taken together, these results reveal how WDR82 can make simultaneous contact with PNUTS and ZC3H4 in the PP1/PNUTS/WDR82/ZC3H4 heterotetramer by using a subset of the contacts used in the PP1/PNUTS/WDR82 and WDR82/ZC3H4 complexes (Figure 2D).

SET1 proteins also bind WDR82, and this interaction antagonizes restrictor function (Hughes et al. 2023). AlphaFold modelling predicted that SETD1A ^60^LQDPR^64^ binds to WDR82 in a similar manner to ZC3H4 ^949^LRDPR^953^, consistent with previous reports (Figures 2D, S1G, H)(Estell et al. 2023; Hughes et al. 2023). A second SETD1A sequence ^74^FSLPVPKF^81^ is predicted to bind the surface of WDR82 bound by the ZC3H4 ^966^VPLSKPSF^971^ and the reversed direction PNUTS ^461^WVCPRPLVL^469^. These sequences structurally align to a consensus sequence ΓxLxxPx_1-2_Ω which we termed LxxP motif and where Γ is a hydrophobic (L/V/F) and Ω an aromatic (F/W) residue (Figure S1F). The predicted interactions of WDR82 with SETD1A and ZC3H4 are thus mutually exclusive.

### PP1 phosphatase stimulates restrictor-mediated termination

To investigate the potential role of PP1/PNUTS in transcription termination by restrictor we expressed the WT and inactive PP1^H66K^-PNUTS (408-940) chimeras and monitored nascent RNA synthesis by Bru-Seq. In this procedure, cells are pulse labelled (20 min) with Bromouridine, then fragmented labelled RNA is immunopurified and RNAseq performed. Bru-Seq signal was normalized to mitochondrial transcripts that act as an internal control. We quantified the ratio of total antisense/sense transcription in the region immediately upstream and downstream of the promoter. Over-expression of PP1^WT^-PNUTS caused a widespread elevation of antisense relative to sense transcription (Figure 3A-C) but the antisense transcripts terminated within a few kb (Figure 3A, B, S2A). (Genes were included in metaplots if the dataset analyzed had at least 1 sense and antisense Bru-Seq reads in the region and will differ between data sets).

**Figure 3.**
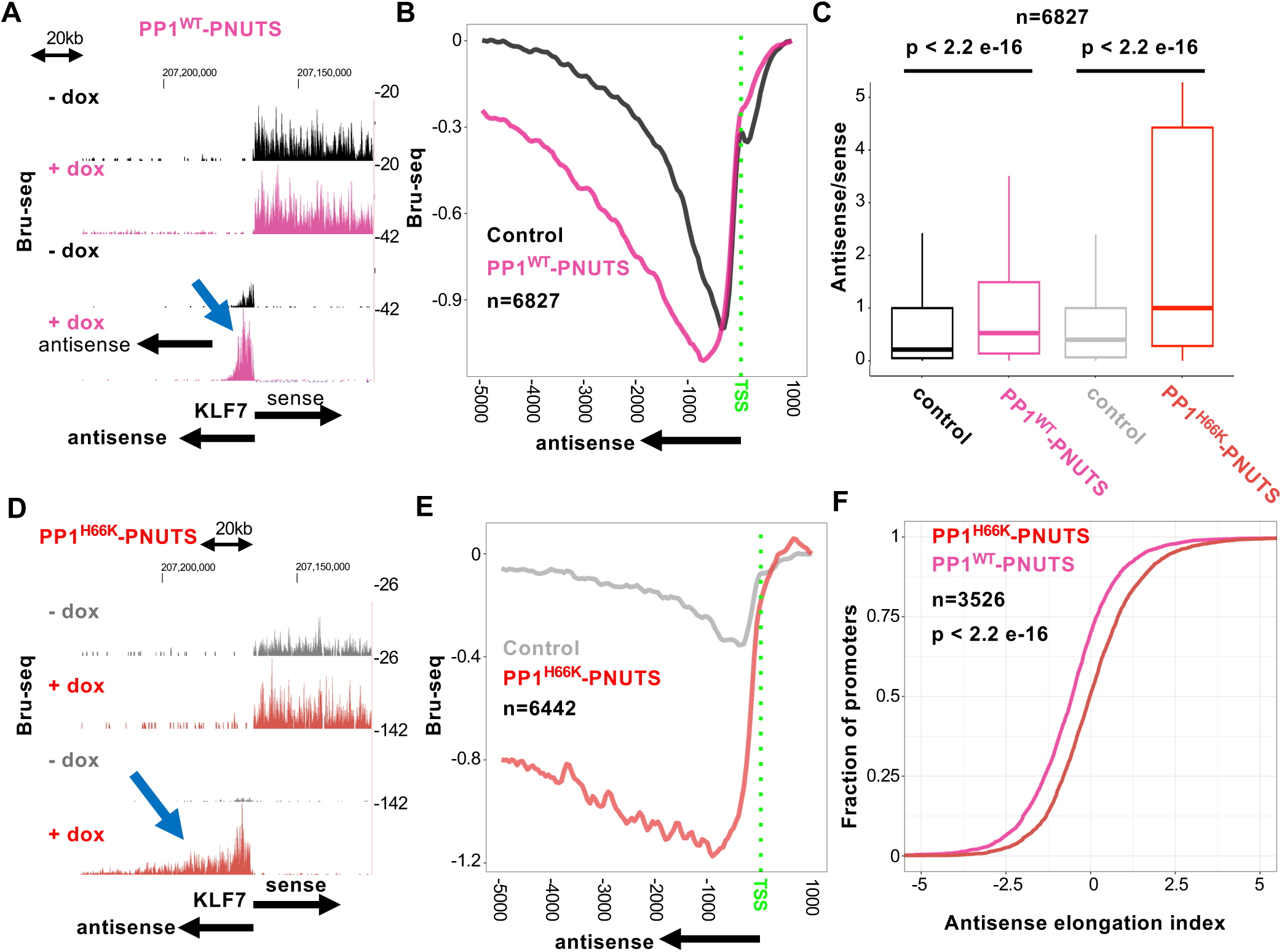
A chimeric PP1^H66K^-PNUTS substrate trap inhibits termination of divergent transcription. **A.** Bru-Seq nascent RNA sequencing at KLF7 for control (-dox) and PP1^WT^-PNUTS (+dox) expressing HEK293 cells. Signals are normalized to mitochondrial reads. Note enhanced divergent antisense transcription (blue arrow) (see Figure S2A). **B.** Metaplots of divergent antisense Bru-Seq signal as in A. Signals are equalized to position +1000. Genes plotted are ≥1kb long and separated from upstream genes by 5kb. **C.** PP1-PNUTS fusion proteins enhance antisense/sense transcription. The ratio of Bru-Seq signals was calculated at each gene for antisense transcripts in the region -1 to -1500 and sense transcripts in the region +1 to +1000 for genes >2kb long and separated by 5kb from other genes. **D.** Bru-Seq nascent RNA sequencing at KLF7 in control (-dox) and PP1^H66K^-PNUTS (+dox) expressing cells. Note enhanced divergent antisense transcription (blue arrow). **E.** Metaplots of divergent antisense Bru-Seq signal as in B for control (-dox) and PP1^H66K^-PNUTS expressing cells +dox). **F.** PP1 inactivation increases the processivity of divergent antisense elongation. Cumulative frequency plots of antisense elongation index (log2 (read density -5000 to -1000)/(read density -1000 to TSS) for PP1-and PP1^H66K^-PNUTS expressing cells.

To evaluate the potential role of protein dephosphorylation on antisense vs sense transcription, we next examined the catalytically inactive PP1^H66K^-PNUTS fusion that acts as a substrate trap. Bru-Seq showed that PP1^H66K^-PNUTS also preferentially enhanced antisense/sense transcription but the effect was more pronounced than for PP1^WT^-PNUTS (Figure 3C) and the length of antisense transcripts were greatly enhanced to often exceed 10kb. This effect was evident at individual genes and some enhancers, and in metaplots of many divergent transcription units (Figure 3D, E, S2B, C). We identified over 2700 genes whose divergent antisense transcription was upregulated by PP1^H66K^-PNUTS at least 2.0 fold in the region between -5000 and the TSS (and separated by 5kb from the nearest upstream gene) (Supplemental Table S2). At over 3000 promoters PP1^H66K^-PNUTS stimulated antisense transcription in the region between -1500 and -1 actually exceeded sense transcription between +1 and +1000 (Figure 3C). To quantify the relative extents of upstream transcription we calculated an antisense elongation index as the log2 ratio of (antisense read density in the region -5000 to -1000)/(antisense read density in the region -1000 to the TSS). Cumulative frequency analysis of these elongation indices showed that antisense transcription elongation is significantly enhanced when the PP1-PNUTS fusion is catalytically inactive (Figure 3F). Together these results show that inactivation of PP1 in the context of a fusion with PNUTS greatly enhances the amount of antisense relative to sense transcription and the distance travelled by antisense directed transcription complexes.

Though antisense transcription is preferentially stimulated, PP1^H66K^-PNUTS also elevated sense transcription to some extent (Figure S2D). PP1^H66K^-PNUTS but not PP1^WT^-PNUTS also elevated transcription in 3’ flanking regions consistent with a role for dephosphorylation by PP1/PNUTS in polyA site dependent transcription termination (Shi et al. 2009; Austenaa et al. 2015; Parua et al. 2018; Cortazar et al. 2019)(Figure S2E, F). Localization of PP1- and PP1^H66K^-PNUTS fusions by ChIP-seq showed that they both had very low level diffuse association along the length of transcribed genes (Figure S3A).

Since PP1^H66K^-PNUTS binds to restrictor (Figure 1E, 2B) we compared its effects on transcription with those of depleting ZC3H4 or ARS2, which aids in restrictor recruitment (Estell et al. 2023; Rouvière et al. 2023). Based on nascent RNA POINT-seq data in HCT116 cells (Estell et al. 2023) we identified 2207 promoters where ZC3H4 degron depletion elevated antisense transcription ≥2X (Figure S3B). Divergent antisense transcription at these ZC3H4-sensitive promoters was strongly upregulated by PP1^H66K^-PNUTS in HEK293 (Figure 4A). In fact PP1^H66K^-PNUTS appeared to have an even more profound effect on divergent transcription at these loci than ZC3H4 depletion (compare Figure 4A with Figure S3B). Similarly transcription units upregulated by ARS2 kd (Rouvière et al. 2023) were also upregulated by PP1^H66K^-PNUTS whereas ARS2 insensitive transcription units were not (Figure 4B, S3C). On the basis of the extensive overlap between the effects of ZC3H4 or ARS2 depletion and PP1^H66K^-PNUTS expression on divergent antisense transcription, we conclude that the PP1^H66K^-PNUTS substrate trap is a potent inhibitor of restrictor-mediated termination. This inhibition of restrictor requires inactivation of PP1 catalytic activity, therefore we conclude that dephosphorylation by PP1 stimulates restrictor function.

**Figure 4.**
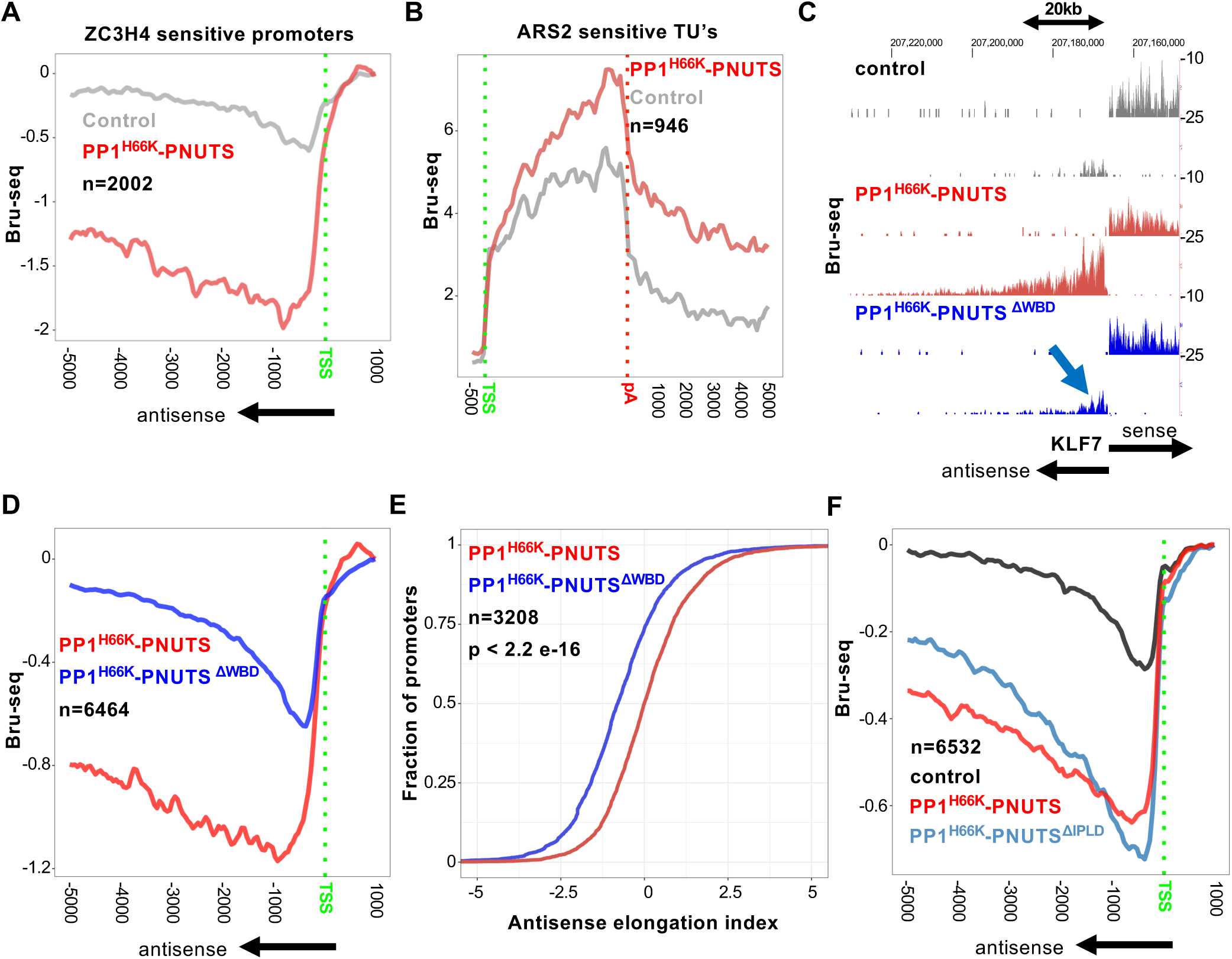
Inhibition of restrictor-mediated termination by PP1^H66K^-PNUTS requires WDR82 binding. **A-B.** Restrictor-mediated termination is inhibited by the PP1^H66K^-PNUTS substrate trap. **A.** Metaplots of Bru-Seq signal in control and PP1^H66K^-PNUTS expressing cells for antisense transcription units upregulated by ZC3H4 degron depletion (see Figure S3B)(Estell et al. 2023). **B.** Metaplots of Bru-Seq signal as in A for transcription units sensitive to depletion of ARS2 (Rouvière et al. 2023). (see Figure S3C). **C. D**, **E.** WDR82 binding is necessary for inhibition of termination by PP1^H66K^-PNUTS. Bru-Seq at KLF7 in control (-dox), PP1^H66K^-PNUTS (replicate 2) and PP1^H66K^-PNUTS^ι1WBD^ expressing cells. Note that deletion of the WDR82 binding domain (WBD) restores termination (blue arrow). **D.** Metaplots of Bru-Seq signal as in C. **E.** Cumulative frequency plots of antisense elongation index for PP1^H66K^-PNUTS and PP1^H66K^-PNUTS^ι1WBD^ expressing cells. Note reduced elongation index for the △WBD mutant. **F.** The PNUTS IPLD motif is not essential for inhibition of termination by PP1^H66K^-PNUTS. Metaplots of Bru-Seq signals for control (-dox), PP1^H66K^-PNUTS (rep 2) and PP1^H66K^-PNUTS^ι1IPLD^ expressing cells. Note inhibition of antisense termination is only slightly impaired by the mutation.

### WDR82 binding is required for inhibition of restrictor termination by PP1^H66K^-PNUTS

The preceding results suggest that PP1 catalytic activity promotes termination by restrictor and that PP1/PNUTS contacts restrictor via WDR82. To test whether phosphatase dead PP1^H66K^-PNUTS must contact restrictor in order to exert its dominant negative effect, we performed nascent RNA sequencing in cells expressing PP1^H66K^-PNUTS^△WBD^ that lacks the WDR82 binding domain (Figure 1A, 2B). Deletion of the WBD effectively abolished the inhibition of restrictor function resulting in a significantly reduced ratio of antisense/sense transcription and antisense elongation index relative to WT (Figure 4C-E, S3D). We conclude that potent inhibition of antisense termination results from recruitment of dominant-negative mutant PP1 to the restrictor via the PNUTS-WDR82 interaction.

According to our AlphaFold modeling of the PPWZ heterotetramer, it is the ZC3H4 ^939^IPLD^942^ motif rather than the identical motif in PNUTS that occupies the cognate binding site on WDR82 (Figure 2C-E). We reasoned that if the PNUTS IPLD motif is dispensable for interaction with restrictor in the PPWZ complex, then mutation of this element would not impair the dominant negative effect of PP1^H66K^-PNUTS on termination. To test this idea, we mutated ^520^IPLD^523^ in PNUTS to GSGS in the context of the PP1^H66K^-PNUTS fusion and monitored nascent RNA synthesis by Bru-Seq. Bru-Seq showed that PP1^H66K^-PNUTS^△IPLD^ inhibited antisense termination only slightly less well than PP1^H66K^-PNUTS^WT^ (Figure 4F, S3E). This result suggests that the IPLD motif in PNUTS is not critical for interaction with WDR82 in agreement with the prediction that in the PPWZ heterotetramer, it is the ZC3H4 IPLD motif that occupies the complementary binding site on WDR82 (Figure 2D, E).

### CTD Ser5 dephosphorylation by PP1/PNUTS and transcription termination

The fact that PP1 phosphatase activity promotes termination by restrictor begs the question of the relevant substrate(s). One candidate is the pol II large subunit (Figure 1E) that co-precipitates with PP1^H66K^-PNUTS and is dephosphorylated at Ser5 residues on its CTD heptad repeats by PP1/PNUTS (Ciurciu et al. 2013; Wu et al. 2018). We tested whether PP1^H66K^-PNUTS affected Ser5 phosphorylation of pol II engaged on genes by ChIP-seq. The results, showed that PP1^H66K^-PNUTS caused Ser5 hyperphosphorylation at the 5’ ends of most genes without a substantial effect on recruitment of total pol II (Figure 5A, B) and this effect was not limited to genes with extensive antisense transcription (Figure 5A, S4A). In contrast to the majority of protein-coding genes, PP1^H66K^-PNUTS did not cause Ser5 hyperphosphorylation at histone or U snRNA genes (Figure 5E, F, S4E, F) that produce short non-adenylated transcripts. Similar results were obtained using an independent anti-Ser5-P monoclonal antibody for ChIP (Figure S4G) and conclusions were unaffected by whether the signal was normalized to the mouse cell spike-in or the snRNA gene internal control (Figures 5A, S4B).

**Figure 5.**
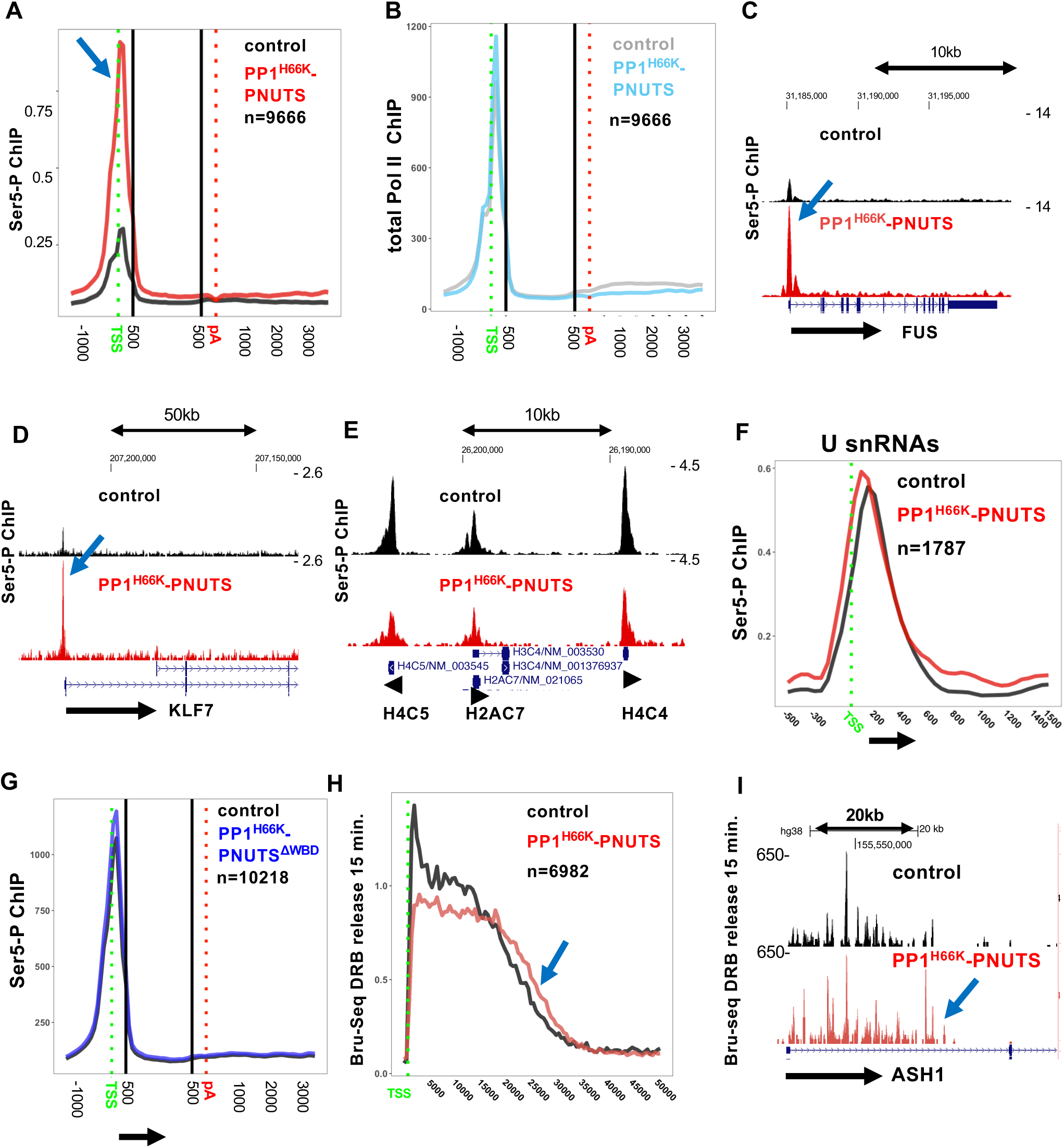
PP1^H66K^-PNUTS causes pol II CTD Ser5 hyperphosphorylation and accelerated transcription. **A.** Pol II Ser5-P ChIP seq metaplots in control (-dox) and PP1^H66K^-PNUTS (+dox) expressing cells. ChIP-signal (3E8 monoclonal anti-Ser5-P) was normalized to a mouse spike-in. Plots were determined for genes longer than 1 kb and separated by 2kb with more than 1 read. Note enhanced Ser5-P in PP1^H66K^-PNUTS cells (blue arrow) (see also Figure S4A, B, G). **B.** Total Pol II ChIP seq metaplots in control (-dox) and PP1^H66K^-PNUTS (+dox) expressing cells. ChIP-signal was normalized to U snRNA genes. Note PP1^H66K^-PNUTS does not alter total pol II recruitment (See Figure S4B). **C, D.** Ser5-P ChIP-seq as in A at the FUS and KLF7 genes showing enhanced phosphorylation induced by PP1^H66K^-PNUTS (blue arrows, see Figure S4C, D). **E**, **F.** PP1^H66K^-PNUTS does not cause Ser5 hyperphosphorylation at histone or U snRNA genes. E, Genome browser screenshot as in C, D at a cluster of histone genes (see Figure S4F). **F**. ChIP seq metaplots as in A of U snRNA genes (see Figure S4E). **G.** WDR82 binding is necessary for inhibition of Ser5-P dephosphorylation by PP1^H66K^-PNUTS. Pol II Ser5-P ChIP seq metaplots in control (-dox) and PP1^H66K^-PNUTS^△WBD^ (+dox) expressing cells. ChIP-signal (monoclonal 3E8) was normalized to U snRNA genes. Note little or no Ser5 hyperphosphorylation (see Figure S4H). **H, I.** PP1^H66K^-PNUTS modestly accelerates transcription. Bru-Seq 15 min after release from a DRB block in control (-dox) and PP1^H66K^-PNUTS (+dox) expressing cells. Read counts in the two data sets were equalized by subsampling in H. Note the wave of transcription travels further with PP1^H66K^-PNUTS (blue arrows) (see Figure S4I, J).

To determine whether WDR82 binding was required for PP1^H66K^-PNUTS to inhibit Ser5 dephosphorylation we conducted anti-Ser5P ChIP before and after inducing expression of PP1^H66K^-PNUTS^△WBD^. This experiment showed no detectable effect of the △WBD mutant on CTD Ser5 phosphorylation (Figure 5G, S4H) consistent with the idea that inhibition of Ser5-P dephosphorylation requires targeting of the substrate trap to the CTD by WDR82 which recognizes phospho-Ser5 heptads (Lee and Skalnik 2008).

To determine whether PP1^H66K^-PNUTS expression and resulting pol II Ser5 hyperphosphorylation affected transcription elongation, we conducted Bru-Seq after release from DRB which blocks elongation at the promoter-proximal pause site (Singh and Padgett 2009). This experiment showed that PP1^H66K^-PNUTS caused the wave of pol II, to advance slightly further into genes 15 minutes after release from the DRB arrest (Figure 5H, I, Figure S4I, J). We could not monitor the speed of divergent antisense transcription in this experiment because it is confounded by the inhibition of termination. In summary, these observations demonstrate a widespread but gene-specific dephosphorylation of CTD Ser5 by PP1/PNUTS at 5’ ends, and that the PP1^H66K^-PNUTS fusion interferes with this process and modestly accelerates transcription. Our results to not exclude the possibility, that dephosphorylation of other PP1/PNUTS substrates is also inhibited by PP1^H66K^-PNUTS including possibly nuclear exosome subunits (Fig. 1F).

### Reduced pausing by CTD Ser5 hyperphosphorylated pol II

To investigate the possibility of a mechanistic connection between pol II Ser5-P dephosphorylation and termination, we monitored transcription by Ser5 hyperphosphorylated pol II and total pol II by eNETseq (Fong et al. 2022). This method can precisely determine the orientation and position of different pol II isoforms by sequencing of nascent RNA fragments protected by the polymerase (Churchman and Weissman 2011; Nojima et al. 2015). eNETseq confirmed that PP1^H66K^-PNUTS caused Ser5 hyperphosphorylation without substantially enhancing total pol II recruitment to promoters. It also confirmed that polymerases engaged in divergent antisense transcription following inhibition of termination by PP1^H66K^-PNUTS are indeed Ser5 phosphorylated (Figure 6A-C, S5A).

**Figure 6.**
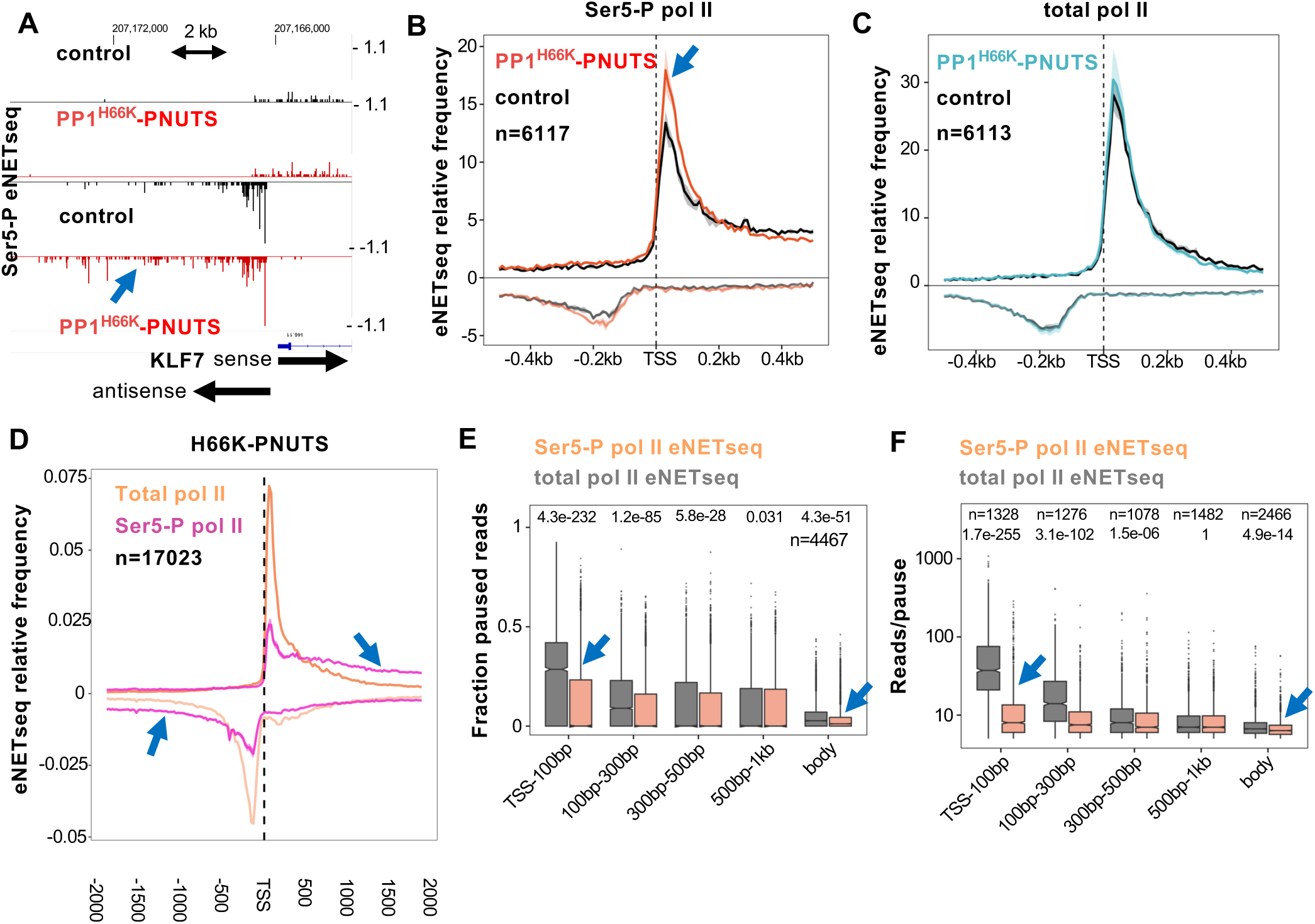
Enhanced elongation by Ser5-P pol II relative to total pol II. **A.** Ser5-P eNETseq reads at KLF7. Note elevated antisense signal (blue arrow) in cells expressing PP1^H66K^-PNUTS (+dox) relative to control (-dox) (see Figure S5A). **B. C.** Metaplots of relative eNETseq signal (2 replicates) for Ser5-P pol II (B) and total pol II (C) in cells expressing PP1^H66K^-PNUTS and control as in A for genes >2 kb long and separated by >5 kb. Subsampling equalized the number of mapped reads in each dataset. Note elevated Ser5-P signal (B, blue arrow) relative to total pol II in cells expressing PP1^H66K^-PNUTS. The center line shows the mean of the replicates and a shaded region shows the minimum and maximum. **D.** Metaplots of relative eNETseq signal (2 replicates) for Ser5-P pol II and total pol II in cells expressing PP1^H66K^-PNUTS (see Figure S5B). Read numbers were equalized by subsampling. Note the relative amount of elongating versus 5’ paused pol II is higher for Ser5-P pol II than total pol II (blue arrows). **E. F.** Reduced pausing by Ser5-P pol II relative to total pol II. Fraction of paused eNETseq reads and reads/pause are plotted for different gene regions. Pauses are positions with >5 reads where the signal is >3 SD above the mean in a 200 bp window. Reads were equalized by subsampling. Data from HEK293 Flp-in cells. (Replicates in Figure S5E, F)

Direct comparison of eNETseq profiles generated from immunoprecipitates of total pol II and the Ser5 phospho-isoform revealed a remarkable difference that has not been emphasized previously and is not obvious from ChIP experiments using the same antibodies. The total pol II profile exhibits the signature peak of promoter-proximal pol II associated with pausing that precedes premature termination or release into productive elongation. Ser5 phosphorylated pol II on the other hand, has a quite distinct profile with relatively less occupancy at promoter-proximal positions and more at distal positions in both the sense and antisense directions (Figure 6D). In other words the Ser5-phosphorylated pol II sub-population behaves as if it transitions from promoter-proximal to promoter-distal positions far better than total pol II. This observation is consistent with previous work showing that inhibition of the Ser5 kinase Cdk7 increased the “pausing index”, a ratio of promoter-proximal: promoter distal pol II density (Ebmeier et al. 2017). Similar results were observed in HEK293 cells +/-dox induction of PP1^H66K^-PNUTS (Figure 6D, S5B) as well as in parental HEK293 (Figure S5C). Examination of the mNETseq data of Nojima et al (2015) revealed a similar contrast between the profiles of Ser5-P pol II and total pol II using independent antibodies for pol II immunopurification (Figure S5D). We investigated the basis for enhanced elongation by Ser5-P pol II by quantifying the fraction of reads at pause sites (pausing propensity) and the number of reads/pause (pause strength) in different regions of the gene. Pause sites are defined as positions with >5 reads where the signal is >3 standard deviations above the mean of the surrounding 200 bp window (Churchman and Weissman 2011; Fong et al. 2022). This analysis revealed that relative to total pol II, the Ser5-P class has much reduced pausing propensity and pause strength both in promoter-proximal regions and within gene bodies (Figure 6E, F, S5E, F). Together these results suggest that the dephosphorylation of CTD Ser5 residues by the PPWZ complex could facilitate transcription termination by increasing pausing and thereby reducing the capacity of pol II to elongate (Figure 7).

**Figure 7.**
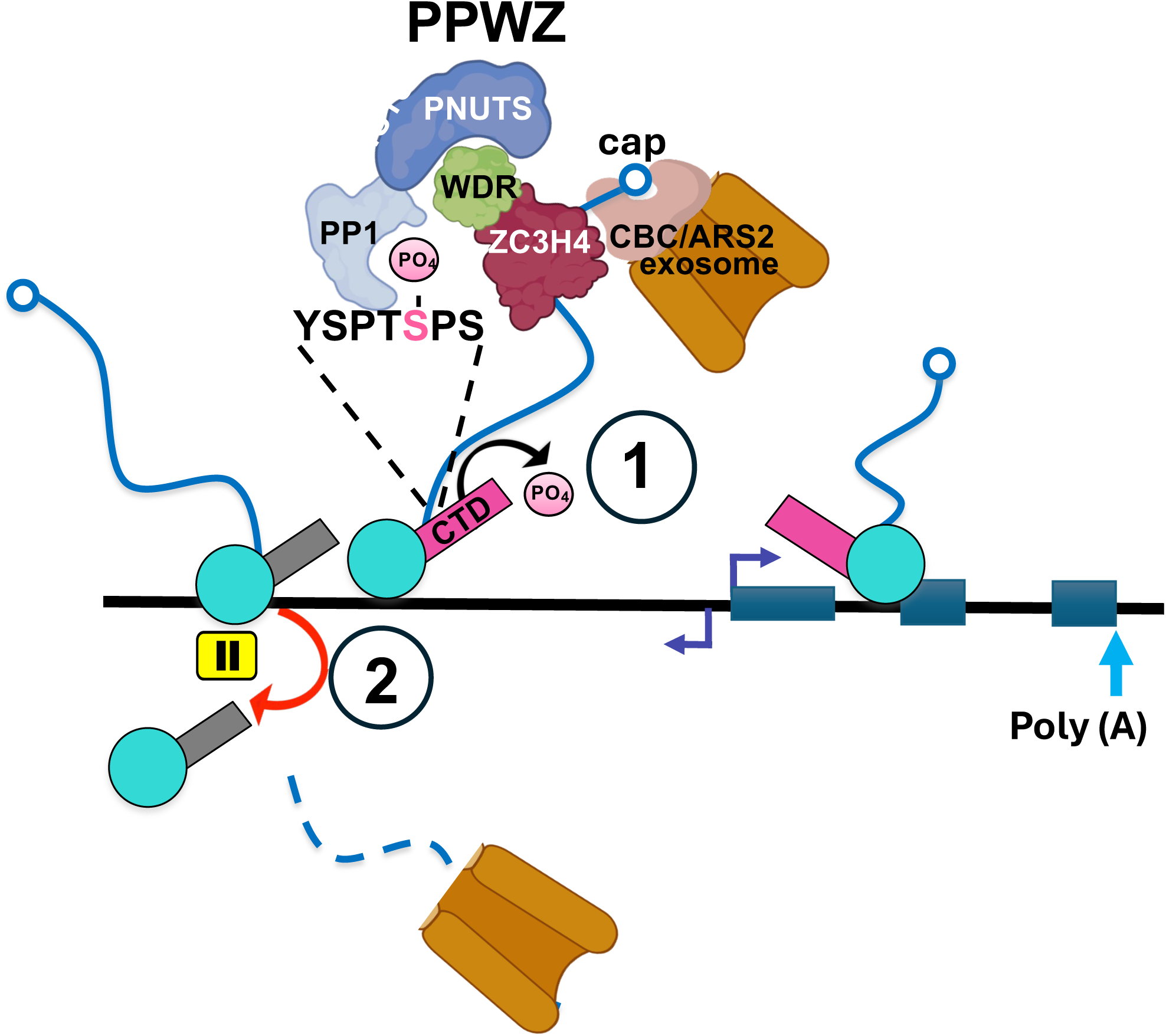
Model of coupled CTD Ser5 dephosphorylation and termination by the PPWZ complex together with the nuclear exosome.

## Discussion

We report here that the dephosphorylation activity of PP1 phosphatase is required for termination of divergent antisense pol II transcription by the restrictor and propose that the physiologically relevant form of restrictor in a heterotetrameric complex of PP1/PNUTS/WDR82/ZC3H4 or PPWZ. The most significant observations in support of these conclusions are: 1. PP1/PNUTS co-purifies with both subunits of restrictor (Figure 1C-E) and binds directly to the WDR82 subunit through a conserved interaction domain in PNUTS (Figure 2A, B). AlphaFold predicts that PP1/PNUTS and WDR82 make multiple contacts with one another that are compatible with ZC3H4 binding to WDR82 in a PPWZ complex (Figure 2D, E). 2. PP1 phosphatase activity stimulates restrictor mediated termination. This conclusion is based on the fact that a substrate trap comprising hypoactive PP1^H66K^ fused to PNUTS strongly inhibits restrictor mediated termination relative to the same fusion with WT PP1 (Figure 3). Inhibition of termination by PP1^H66K^-PNUTS requires the WDR82 binding domain or WBD through which it contacts the restrictor (Figure 2B, 4C-E). Recruitment of inactive, but not active PP1 to restrictor, therefore works as a dominant negative inhibitor of termination. Stimulation of antisense transcription by PP1^WT^-PNUTS (Figure 3A, B,) is also likely caused by binding to the WDR82 subunit of restrictor, but without inhibition of PP1, it has a relatively small effect. 3. CTD Ser5 dephosphorylation by PPWZ contributes to termination by restrictor. This conclusion is suggested by the fact that PP1^H66K^-PNUTS expression hyperphosphorylates CTD Ser5 and inhibits termination, and both activities are abolished by deleting the PNUTS WBD (Figs. 4C-E, 5G). The importance of the PNUTS-WDR82 interaction for localization of PP1 to the restrictor suggests that the presence of WDR82 in other protein complexes may also serve to recruit this phosphatase.

We propose a functional coupling between termination and CTD Ser5 dephosphorylation by the PPWZ complex (Figure 7). Notably WDR82 binds specifically to Ser5 phosphorylated CTD heptads (Lee and Skalnik 2008) and both WDR82 and ZC3H4 associate with Ser5 (TFIIH) phosphorylated and not unphosphorylated CTD (Ebmeier et al. 2017). Ser5-P binding by WDR82 and RNA binding by ZC3H4 could help to direct the phosphatase to divergently transcribing pol II complexes (Murray et al. 1997; Lee and Skalnik 2008). A rationale for how Ser5 dephosphorylation facilitates termination is suggested by our observation that the frequency and strength of transcriptional pausing is much reduced for the Ser5 hyperphosphorylated isoform relative to total pol II (Figure 6D-F). Dephosphorylation of Ser5-P by PPWZ would convert pol II from a faster low pausing to a slower high pausing state that is presumably more susceptible to the restrictor since pausing is a universal pre-condition for termination (Ray-Soni et al. 2016). Our models of PPWZ predict that a helix of PNUTS can occupy the PP1 active site and mimic the interactions made by PP1 inhibitor-2 (Hurley et al. 2007) (Figure 2C). While the functional significance of such autoinhibition is unknown, it is interesting to note that in the integrator, another CTD Ser5 phosphatase/terminator complex, an inhibitory loop of INTS6 occupies the PP2A active site (Fianu et al. 2024).

Additional potential substrates of PP1/PNUTS that may function in restrictor mediated termination include nuclear exosome associated factors that are implicated in this process by virtue of their proximity to ZC3H4 (Estell et al. 2023; Rouvière et al. 2023). We found that MTREX, ZC3H18, ZCCHC8, EXOSC3, 6 and 8 were specifically enriched, like pol II large subunit in IP’s of PP1^H66K^-PNUTS relative to PP1^WT^-PNUTS (Figure 1F). While fusions of PP1 with interacting proteins (PIPs) were first developed as substrate binding traps (Wu et al. 2018) our work shows that inducible expression of WT and mutant PP1-PIP fusions can reveal new functions of specific PP1 holoenzymes which have been difficult to disentangle previously (Ferreira et al. 2019).

This work highlights two emerging principles in control of pol II termination: the involvement of CTD Ser5-P and the function of phosphatases in this final step of the transcription cycle. Ser5-P recognition is central to premature termination of yeast transcription by the Nrd1-Nab3-Sen1 (NNS) pathway. Nrd1 specifically binds Ser5-P CTD heptads (Vasiljeva et al. 2008) and works together with the RNA binding protein Nab3 and the helicase Sen1 to terminate short non-coding transcripts (Porrua and Libri 2015). The yeast NNS complex and the mammalian PPWZ complex proposed here have a number of features in common:1) they both bind phospho-Ser5-P pol II, 2) they bind the nascent RNA, and 3) they interact with the nuclear exosome (Vasiljeva and Buratowski 2006; Estell et al. 2023; Rouvière et al. 2023)(Figure 1F).

The integral function of PP1/PNUTS in restrictor mediated termination reported here adds to previous studies suggesting an ancient role of this phosphatase in pol II termination. PP1/PNUTS functions in polyA site dependent termination in organisms ranging from mammals to yeasts and in J base-dependent termination in trypanosomes (Parua et al. 2018; Sanchez et al. 2018; Cortazar et al. 2019; Carminati et al. 2023; Kieft et al. 2023). In addition the PP2A phosphatase activity of the integrator complex, which also dephosphorylates CTD Ser5, promotes premature termination near the 5’ ends of genes (Huang et al. 2020; Zheng et al. 2020; Fujiwara et al. 2023). The proposal that CTD Ser5 dephosphorylation may facilitate termination by conversion of pol II to a high pausing state could therefore apply to multiple termination mechanisms not only upstream of genes but also downstream and within genes.

### Limitations

We report experimental evidence for several protein:protein interactions within the PPWZ complex and predicted structures at some interfaces, but we have not determined the complete composition or subunit stoichiometry of the complex, nor have we definitively determined its in vivo substrate specificity which is still very challenging for individual PP1 complexes. The model that CTD Ser5 dephosphorylation is necessary for restrictor mediated termination is consistent with the available data, but remains a hypothesis to be tested further.

How Ser5-P phosphorylation affects transcriptional pausing possibly by modulating pol II association with positive and elongation factors remains unknown.

**Author contributions:** R.T and D.B. conceived the project. R.F. S.M. K.H., R.T. and D.B. designed experiments. B.E. and N.F. performed all experiments and B.E. and R.S. performed bioinformatics. A.S. performed Mass Spectrometry, R.F performed AlphaFold modelling. R.F., R.T., K.H., and D.B. wrote the paper with input from all authors.

## Acknowledgements

We thank G. Natoli for helpful discussions and J. Rouviere and T. Jensen for ARS2 sensitive gene lists. R.S is supported by the UC Denver RNA Bioscience Initiative. This work was supported by the UC Denver RNA Bioscience Initiative and NIH grant R35GM118051 to D.B. University of Colorado Mass Spectrometry Shared Resource Facility is supported by the National Cancer Institute through the Cancer Center Support Grant (P30CA06934). This work was supported by the Francis Crick Institute which receives its core funding from Cancer Research UK (CC2102), the UK Medical Research Council (CC2102), and the Wellcome Trust (CC2102). This research was funded in whole, or in part, by the Wellcome Trust CC2102. For the purpose of Open Access, the author has applied a CC BY public copyright license to any Author Accepted Manuscript version arising from this submission.*·*

## Materials and Methods Human cell lines

**HEK293 cells** were infected with lentiviruses based on pCW57 a gift of Adam Karpf (Addgene 89180) and selected with blasticidin (10 μg/ml) for doxycycline inducible expression of PNUTS and PP1-PNUTS fusions. Doxycycline induction (2 µg/mL) was for ∼24 hr.

## Immunoprecipitation

Nuclear extracts were prepared from induced lentiviral transduced cells (24 hr., 2 μg/ml doxycycline) by a modification of the procedure of Baluapuri et al (Baluapuri et al. 2019). Cells were lysed in 10 mM HEPES pH 7.9, 0.34 M sucrose, 3 mM CaCl_2_, 2 mM magnesium acetate, 0.1 mM EDTA, 0.5% NP-40, nuclei pelleted and benzonase digested (200 U/ml) for 40 min at 4° with nutating in 20 mM HEPES pH 7.9, 0.1 mM EDTA, 10% glycerol, 150 mM potassium acetate, 1.5 mM MgCl2,. An equal volume of 20 mM HEPES pH 7.9, 0.1 mM EDTA, 10% glycerol, 600 mM NaCl, 1.5 mM MgCl2 was added and nuclei were nutated 30 min. at 4°. Insoluble material was pelleted in a microfuge 15,000 rpm 15 min and the supernatant was diluted with an equal volume of 2% v/v Nonidet P-40 1% w/v deoxycholate, 0.2% w/v SDS, 50 mM Tris pH 8, 5 mM EDTA.

Nuclear extract (1-2 mg) was immunoprecipitated with rabbit polyclonal anti Avitag antibody (2 hr, 4°) and collected on Protein A Dynabeads, washed 3X rapidly with 50 mM Tris-Cl, pH 8.0, 150 mM NaCl, 0.5% NP-40 and eluted with 1% Formic Acid and 28% Acetonitrile, for Mass spectrometry or Western blotting.

### Immunoblotting

Immunoblots were developed with HRP conjugated swine anti-rabbit secondary antibody (DAKO, P0217) and ECL Plus (Perkin Elmer, NEL103E001EA).

### Mass spectrometry

Samples (4-6 replicates) were reduced, alkylated, and digested using S-Trap^TM^ micro filters (Protifi, Huntington, NY) according to the manufacturer’s protocol. Digested peptides were cleaned using Pierce^TM^ C18 Spin Tips (Thermo Scientific), dried in a vacuum centrifuge, and resuspended in 0.1% FA in mass spectrometry-grade water. Peptides were loaded into autosampler vials and analyzed directly using a NanoElute liquid chromatography system (Bruker, Germany) coupled with a timsTOF SCP mass spectrometer (Bruker, Germany). Peptides were separated on a 75 µm i.d. × 15 cm separation column packed with 1.9 µm C18 beads (Bruker, Germany) over a 90-minute elution gradient. Buffer A was 0.1% FA in water and buffer B was 0.1% FA in acetonitrile. Instrument control and data acquisition were performed using Compass Hystar (version 6.0) with the timsTOF SCP operating in parallel accumulation-serial fragmentation (PASEF) mode under the following settings: mass range 100-1700 m/z, 1/k/0 Start 0.7 V s cm^-2^ End 1.3 V s cm^-2^; ramp accumulation times were 166 ms; capillary voltage was 4500 V, dry gas 8.0 L min^-1^ and dry temp 200°C. The PASEF settings were: 5 MS/MS scans (total cycle time, 1.03 s); charge range 0–5; active exclusion for 0.2 min; scheduling target intensity 20,000; intensity threshold 500; collision-induced dissociation energy 10 eV.

Fragmentation spectra were searched against the UniProt human proteome database using the MSFragger-based FragPipe computational platform (Kong et al. 2017). Contaminants and reverse decoys were added to the database automatically. The precursor-ion mass tolerance and fragment-ion mass tolerance were set to 15 and 20 ppm, respectively. Fixed modifications were set as carbamidomethyl (C), and variable modifications were set as oxidation (M), two missed tryptic cleavages were allowed, and the protein-level false discovery rate (FDR) was ≤ 1%.

### Bru-Seq

Nascent RNA sequencing by Bru-Seq was performed as described (Paulsen et al. 2014; Sheridan et al. 2019) with minor modifications. HEK293 cells + and – doxycycline induction (2 μg/ml, 24hrs) were incubated with 2 mM Bromouridine in fresh medium + or – doxycycline for 10 or 20 min. Bru-Seq following release from a DRB block (Sheridan et al. 2019) was carried out by addition of BrU (2mM) 10 minutes before harvesting. Labeled RNA (50 μg) was fragmented with ZnCl2 (10mM, 70° 8 min, stopped with 10 mM EDTA) and immunopurified in PBS, 0.05% Triton X-100, 1 mM DTT, + RNase inhibitor for 1 hr at 4° using 3D4 monoclonal anti-BrdU (3 μg) immobilized on protein G Dynabeads. Beads were washed 3X in the same buffer and RNA was eluted in 5mM DTT at 95° for 3 min. Eluted RNA was rRNA depleted (Baldwin et al. 2021) and RNA-seq libraries were made with the KAPA RNA HyperPrep kit. Sequencing was on an Illumina NovaSeq 6000 (2x150). Reads were mapped to the hg38 UCSC human genome with Bowtie2 version 2.3.2. PCR duplicates were removed using bbtools clumpify and adapters were trimmed using bbtools bbkuk version 39.01. After filtering out rRNA, reads were mapped to the hg38 UCSC human genome with Bowtie2 version 2.3.2.

### Bru-Seq analysis

Metaplots include all genes longer than 1 kb, separated by >5 kb, containing at least one read in the region, and were in common between samples plotted. Read counts were normalized to total counts on the mitochondrial chromosome. Antisense/Sense boxplots for antisense signal -5kb-TSS / sense signal TSS-+1.0 kb genes >2kb long and separated by 5kb from other genes. Antisense elongation index is the log2 (antisense read density in the region -5000 to - 1000)/(antisense read density in the region -1000 to the TSS).

### AlphaFold modelling

AlphaFold 2.3.0 with full sequence databases was used to model structure predictions with the multimer setting enabled (Evans et al. 2022). Full-length proteins fasta sequences from Uniprot.org were used as input unless otherwise indicated. Default five seeds was used per each five model parameters, 25 in total. Top ranked prediction as indicated by pLDDT was used in the study (unless otherwise indicated).

### ChIP-seq

ChIP was performed as described (Glover-Cutter et al. 2008; Cortazar et al. 2022). Cross-linked human cell pellets were spiked with cross-linked mouse cells prior to sonication with a Covaris ME220. 10 μl of rabbit anti-pan pol II CTD serum 5 μg anti-Avitag antibody, 4 μg of 3E8 rat monoclonal anti-Ser5-P CTD together with rabbit anti-Rat IgG or 5 μg of 4H8 mouse monoclonal anti-Ser5-P CTD was used for each IP with 1 mg of cross-linked extract. Libraries were sequenced on an Illumina NovaSeq 6000 (2x150). Reads were mapped to the hg38 UCSC human genome with Bowtie2 version 2.3.2. PCR duplicates were removed using bbtools clumpify and adapters were trimmed using bbtools bbkuk version 39.01 Read coordinates were collapsed and centered using deeptools bamCoverage version 3.5.1.

Pol II ChIP-seq, metaplots include all genes longer than 1 kb, separated by >2 kb, containing at least one read in the region, and were in common between all ChIP-seq. In some cases read counts were normalized to signal over snRNA genes.

### Point-seq analysis

Reads were mapped to the hg38 UCSC human genome with Hisat version 2.1.0. PCR duplicates were removed using bbtools clumpify and adapters were trimmed using bbtools bbkuk version 39.01 Bigwigs were made using deeptools bamCoverage version 3.5.1. filtering out rRNA, normalizing RPKM.

Metaplots include all genes longer than 1 kb, separated by >2 kb, containing at least one read in the region, and were in common between all samples. To calculate the relative signal for each gene, the signal for each bin was divided by the total signal within the region plotted. The mean relative signal was then calculated by averaging the relative signal across the genes plotted for each individual replicate. The replicates were then averaged with the shaded region representing the min and max range for the three replicates, normalized to the +1000 position. Adding the replicate signals together and selecting all genes longer than 1 kb, separated by >5 kb, containing at least one read in the antisense region 5kb to 0.5kb upstream of the TSS. We then selected genes that had a greater than 2-fold increase in the antisense signal - TMP/+TMP.

### eNET-seq

eNET-seq experiments employed optimized MNase (NEB) digestion conditions, RNA decapping with recombinant *S. pombe* Dcp1-Edc1-Dcp2, and incorporation of 12 base UMIs in library construction (Fong et al. 2022). Pol II immunoprecipitation was with polyclonal rabbit anti pan-pol II CTD or rat monoclonal anti-Ser5-P CTD 3E8 (Chromotek) with rabbit anti-rat secondary antibody.

### eNET-seq read processing

eNET-seq libraries were made with the QIAseq miRNA UDI Library kit using 12 base UMI’s and sequenced on an Illumina NovaSeq 6000 (2x150). Adapters were trimmed using cutadapt (v2.3) and reads were aligned to the hg38 human genome using Bowtie2 (v2.3.2). PCR duplicates were removed using UMI-tools (v0.5.4) and read coordinates were collapsed to a single base pair coordinate corresponding to the RNA 3’ end. Reads were filtered to only include those with a mapping quality score >=10 and to remove reads aligning to snoRNA genes. To generate UCSC genome browser tracks, BigWig coverage files were generated from the aligned/filtered reads. For eNET-seq 5’ metaplots and pausing analysis, reads were further filtered to remove those that did not align within 5 kb of a protein coding gene. Metaplots include all genes longer than 2 kb and separated by >5 kb. In addition, genes had to contain at least one read and the bottom 10% of genes with the lowest signal were excluded. Each 5’ region was divided into 10 bp bins. Read counts in each bin were normalized by library size and the size of the bin. The relative signal (relative frequency) was calculated separately for each gene by dividing the signal in each bin by the sum of the signal for the entire plotted region. The relative signal was then averaged for each bin. For Fig 6B-C and S5C these values were multiplied by 1000 to remove small decimals. The mean relative signal was calculated separately for each biological replicate. The shaded region shows the minimum and maximum mean signal for the replicates and the center line shows the mean for the replicates.

### eNET-seq read subsampling

eNET-seq read subsampling was performed as previously described (Fong et al. 2022) To compare pol II pausing for different TSS regions (Fig 6E,F, S5E,F), reads were subsampled so each gene had the same number of aligned/filtered reads for all datasets used in the comparison. Subsampling was performed separately for each individual gene and involved identifying the dataset with the fewest reads for each gene. Reads aligning to the gene were then randomly subsampled for the remaining datasets so all samples used in the comparison had the same number of reads for the gene. To compare pol II pausing for gene body regions (Fig 6E,F, S5E,F), reads were subsampled for each gene body region (+500 bp – pAS) as described above.

### Quantification of pol II pausing

Pause sites were located as previously described (Fong et al. 2022). Pauses were located by identifying positions where eNET-seq signal was >3 standard deviations above the mean for the surrounding 100 bp on each side, there were at least 5 reads at the pause site, and at least 5 additional reads within the window. Pause sites identified within 2 bp of 5’ and 3’ splice sites were excluded from downstream analysis. The fraction of pause reads was calculated by dividing the total number of reads aligned to pause sites by the total number of reads for the region. Genes were filtered to only include those >2 kb long, separated by >5 kb, and with at least 1 pause in TSS regions (TSS - +1 kb) and a total read density of at least 1 read/kb within the gene body region (+1 kb – pAS). To assess pause strength, the average number of reads/pause was calculated for genes filtered as described above. In addition, for each region genes were further filtered to only include those with at least 1 pause site for the region.

### DATA AVAILABILITY

Sequencing datasets are deposited in GEO GSE269357, GSE269358), GSE269359.

