## Supplemental Figs S1-S5 for "PP1 PNUTS binds the “restrictor” and dephosphorylates RNA pol II CTD Ser5 to stimulate transcription termination"

### Figure S1

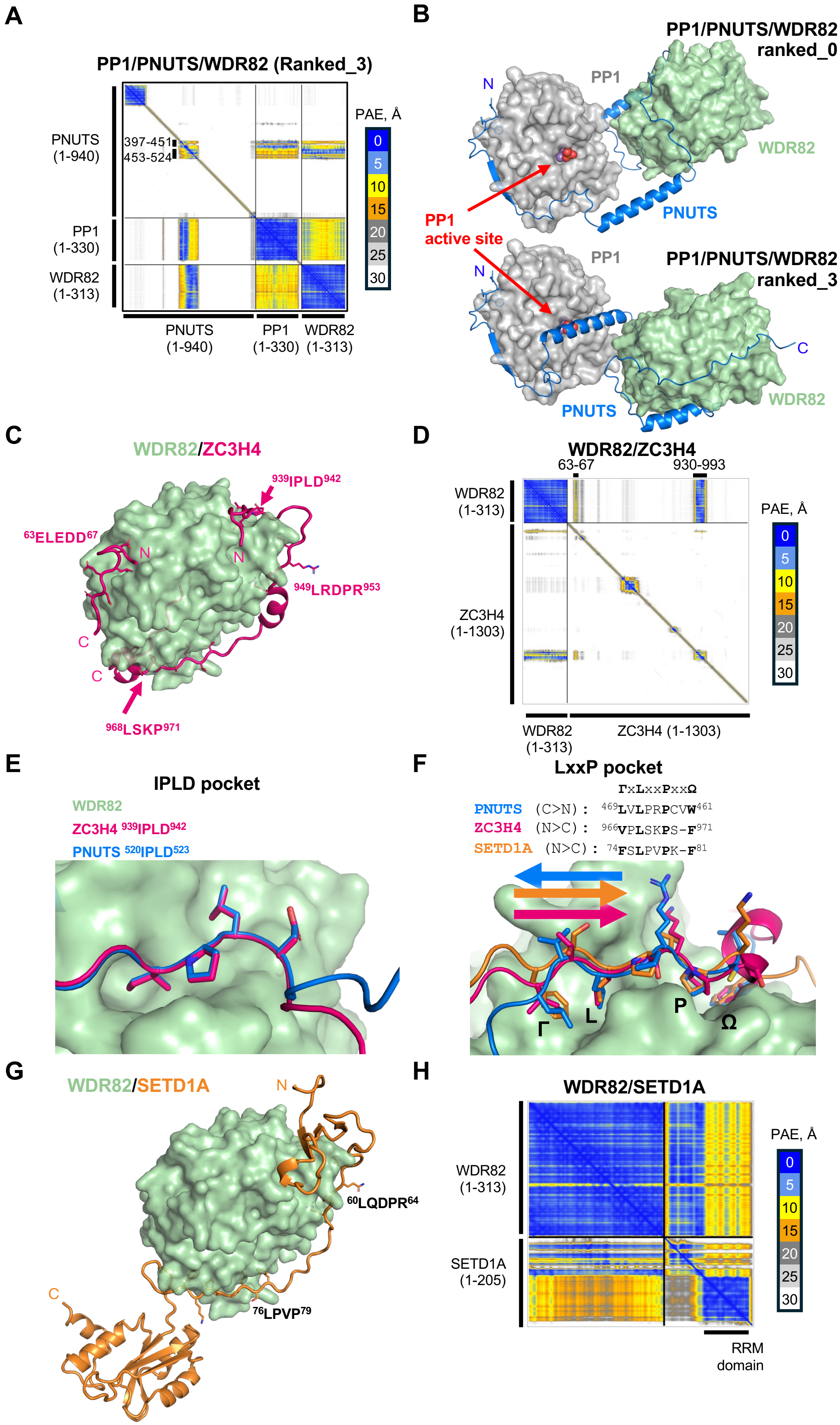

A
PP1/PNUTS/WDR82 (Ranked_3)
PNUTS (1-940)
397-451
453-524
PP1 (1-330)
WDR82 (1-313)
PNUTS (1-940)
PP1 (1-330)
WDR82 (1-313)
PAE, Å
0
5
10
15
20
25
30
B
PP1/PNUTS/WDR82 ranked_0
PP1
N
WDR82
PNUTS
PP1 active site
PP1
PP1/PNUTS/WDR82 ranked_3
N
C
PNUTS
WDR82
C
WDR82/ZC3H4
939IPLD942
63ELEDD67
N
N
949LRDPR953
C
C
968LSKP971
D
WDR82/ZC3H4
63-67
930-993
WDR82 (1-313)
ZC3H4 (1-1303)
PAE, Å
0
5
10
15
20
25
30
WDR82 (1-313)
ZC3H4 (1-1303)
E
IPLD pocket
WDR82
ZC3H4 939IPLD942
PNUTS 520IPLD523
F
LxxP pocket
ΓxLxxPxxΩ
PNUTS (C>N): 469LVLPRPCVW461
ZC3H4 (N>C): 966VPLSKPS-F971
SETD1A (N>C): 74FSLPVPK-F81
Γ
L
P
Ω
G
WDR82/SETD1A
N
60LQDPR64
C
76LPVP79
H
WDR82/SETD1A
WDR82 (1-313)
SETD1A (1-205)
PAE, Å
0
5
10
15
20
25
30
RRM domain

**Supplemental Figure Legends**

**Supplemental Figure S1. AlphaFold predictions of WDR82 interactions with PNUTS, ZC3H4 and SET1.**

**A.** Predicted aligned error plot of the AlphaFold prediction of a ternary complex of PP1/PNUTS with WDR82 shown in Figure 2C where a helix of PNUTS (429-452) occupies the active site of PP1 and the WBD region of PNUTS (453-524) binds WDR82. PP1/PNUTS and PNUTS/WDR82 interactions are separated by a very short but flexible linker, that results in low correlation scores by PAE for the positions of the relevant residues in WDR82 and PP1.

**B.** Comparison of alternative AlphaFold predictions of a ternary complex of PP1/PNUTS with WDR82 in which a helix of PNUTS (red arrows) is either unbound (top 6/25 models) and the PP1 active site is exposed, or this helix occupies the PP1 active site (bottom, 19/25 models) similar to the complex of PP1 inhibitor 2 with PP1 [44].

**C**, **D.** AlphaFold prediction and predicted aligned error plot of the complex between WDR82 and ZC3H4. SLIMs $^{63}$ELEDD$^{67}$, $^{939}$IPLD$^{942}$, $^{949}$LRDPR$^{953}$ and $^{966}$VPLSKPSF$^{971}$ of ZC3H4 contact WDR82, two of which ($^{939}$IPLD$^{942}$,$^{949}$LRDPR$^{953}$) are predicted to persist in the tetrameric PPWZ complex.

**E.** Predicted competition between identical IPLD motifs in PNUTS and ZC3H4 for binding to WDR82.

**F.** Predicted competition between LxxP motifs in PNUTS, ZC3H4 and SETD1A for binding to WDR82.

**G, H.** AlphaFold prediction and the predicted aligned error plot of the SETD1A/WDR82 complex. SLIMs in SETD1A that interact with WDR82 are marked.

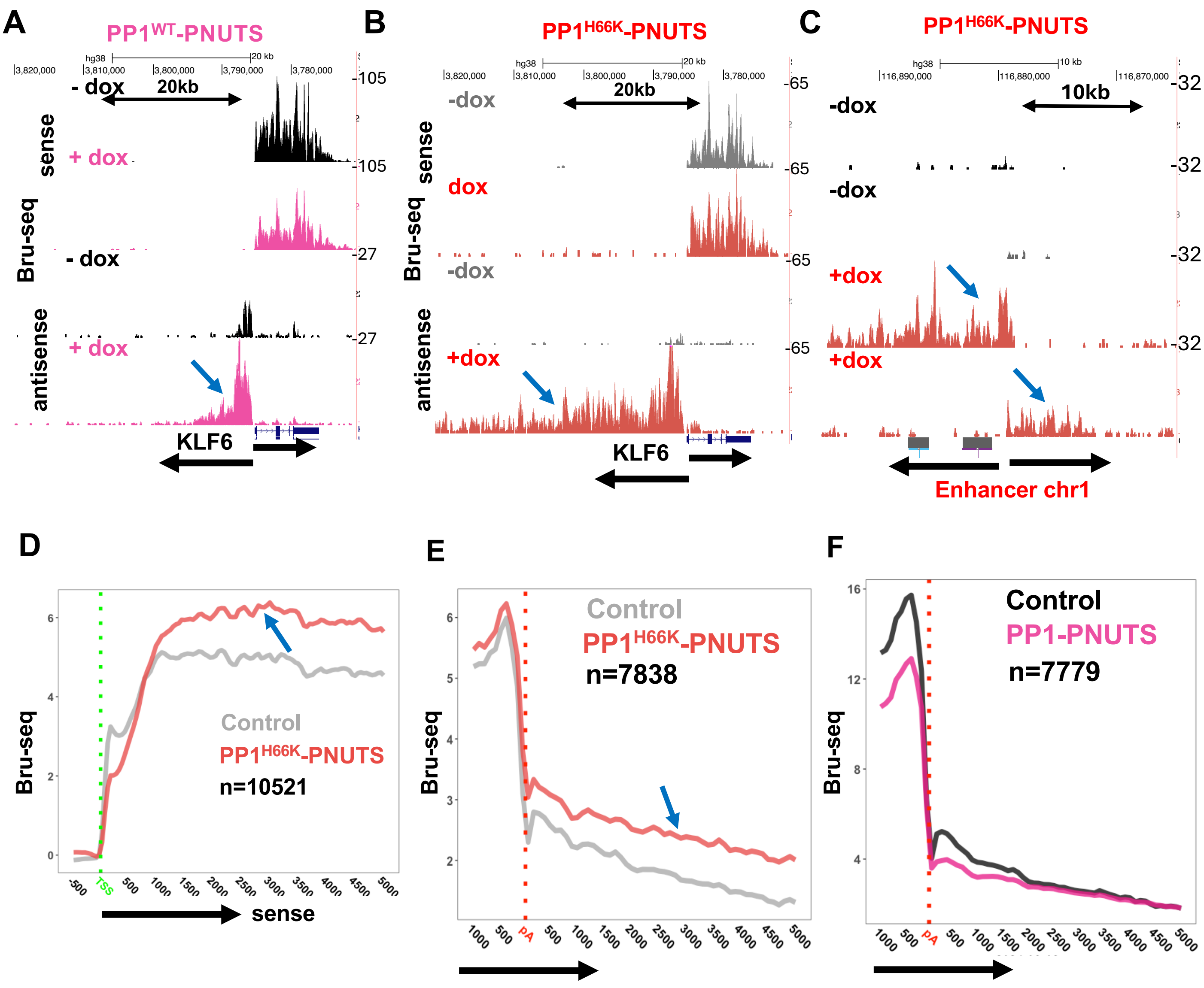
A
PP1WT-PNUTS
- dox
20kb
sense
+ dox
Bru-seq
- dox
antisense
+ dox
KLF6
105
105
27
27
B
PP1H66K-PNUTS
-dox
20kb
sense
dox
Bru-seq
-dox
antisense
+dox
KLF6
65
65
65
65
C
PP1H66K-PNUTS
-dox
10kb
-dox
+dox
+dox
Enhancer chr1
32
32
32
32
D
Bru-seq
Control
PP1H66K-PNUTS
n=10521
TSS
sense
E
Bru-seq
Control
PP1H66K-PNUTS
n=7838
pA
F
Bru-seq
Control
PP1-PNUTS
n=7779
pA

Figure S2

**Supplemental Figure S2** Related to Figure 3

**A chimeric PP1-PNUTS substrate trap blocks termination of divergent transcription**

**A.** Bru-Seq nascent RNA sequencing at KLF6 for control (-dox) and $PP1^{WT}$-PNUTS (+dox) expressing HEK293 cells. Note enhanced divergent antisense transcription (blue arrow) that terminates earlier than that induced by $PP1^{H66K}$-PNUTS (see Figure S2B).

**B.** Bru-Seq nascent RNA sequencing at KLF6 in control (-dox) and $PP1^{H66K}$-PNUTS (+dox) expressing HEK293 cells as in A. Note enhanced divergent antisense transcription (blue arrow).

**C.** Bru-Seq nascent RNA sequencing as in B of an enhancer region with elevated bidirectional transcription (blue arrows).

**D.** Metaplots of sense Bru-Seq signal for control (-dox) and $PP1^{H66K}$-PNUTS expressing cells (+dox). Note sense transcription is elevated (blue arrow) but by less than antisense transcription (see Figure 3C, E).

**E, F.** Inhibition of termination downstream of polyA sites by $PP1^{H66K}$-PNUTS but not $PP1^{WT}$-PNUTS. Metaplots of Bru-Seq signal. Note enhanced signal in the termination zone downstream of polyA sites specifically when $PP1^{H66K}$-PNUTS is expressed (blue arrow).

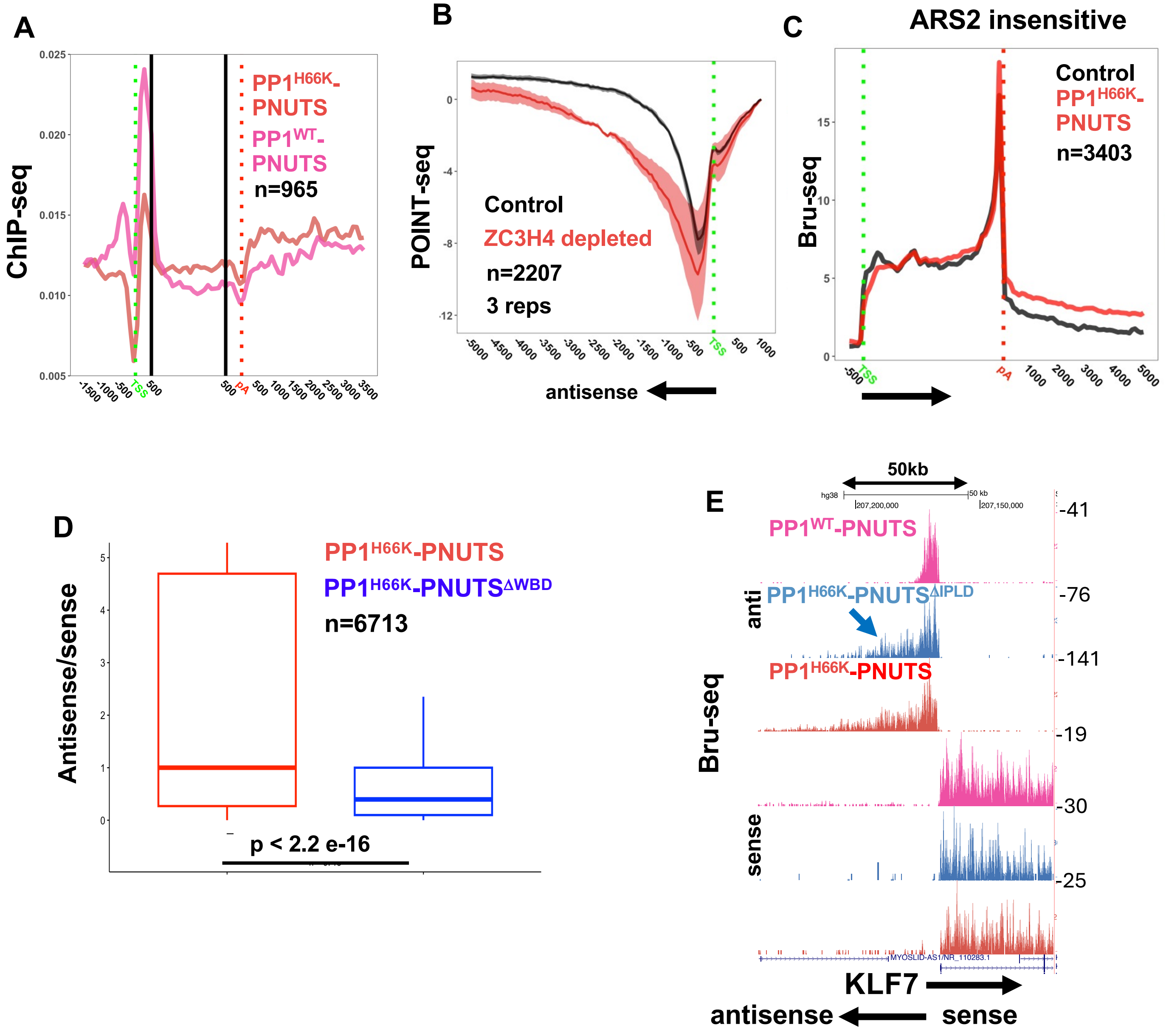
A
ChIP-seq
PP1H66K-PNUTS
PP1WT-PNUTS
n=965
TSS
pA
B
POINT-seq
Control
ZC3H4 depleted
n=2207
3 reps
TSS
antisense
C
ARS2 insensitive
Bru-seq
Control
PP1H66K-PNUTS
n=3403
TSS
pA
D
Antisense/sense
PP1H66K-PNUTS
PP1H66K-PNUTSΔWBD
n=6713
p < 2.2 e-16
E
50kb
Bru-seq
anti
sense
PP1WT-PNUTS
PP1H66K-PNUTSΔIPLD
PP1H66K-PNUTS
-41
-76
-141
-19
-30
-25
KLF7
antisense
sense

Figure S3

**Supplemental Figure S3** Related to Figure 4

**A.** ChIP-seq of Avitag PP1- and PP1$^{H66K}$-PNUTS shows low level diffuse distribution over transcribed genes. Genes shown in the top 10% of transcribed genes are shown,

**B.** Nascent RNA POINT-seq for 2207 divergent antisense transcription units that are 2X upregulated by ZC3H4 degron depletion in HCT116 cells (data from [8]).

**C.** Metaplots of Bru-Seq signal in PP1$^{H66K}$-PNUTS and control (-dox) cells for transcription units insensitive to depletion of ARS2 [9].

**D.** Enhanced antisense/sense transcription induced by PP1$^{H66K}$-PNUTS requires the PNUTS WDR82 binding domain (WBD). The ratio of antisense/sense Bru-Seq signals was calculated as in Figure 3C.

**E**. Bru-Seq nascent RNA sequencing at KLF7. Note divergent antisense transcription in PP1$^{H66K}$-PNUTS$^{\Delta IPLD}$ (blue arrow) is almost as extensive as in PP1$^{H66K}$-PNUTS.

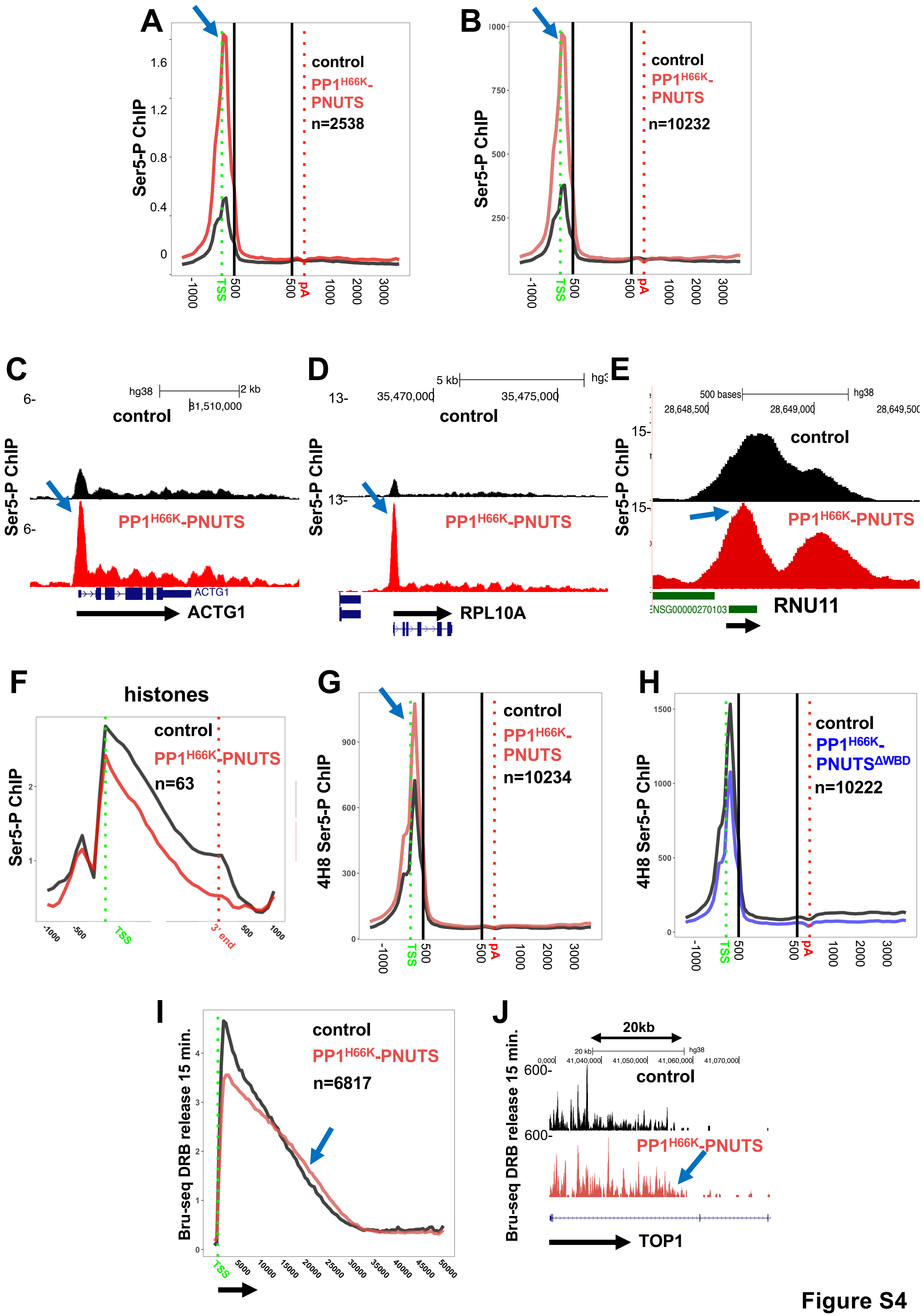
A
Ser5-P ChIP
control
PP1H66K-PNUTS
n=2538
TSS
pA
B
Ser5-P ChIP
control
PP1H66K-PNUTS
n=10232
C
Ser5-P ChIP
control
PP1H66K-PNUTS
ACTG1
D
control
PP1H66K-PNUTS
RPL10A
E
control
PP1H66K-PNUTS
RNU11
F
histones
Ser5-P ChIP
control
PP1H66K-PNUTS
n=63
G
4H8 Ser5-P ChIP
control
PP1H66K-PNUTS
n=10234
H
4H8 Ser5-P ChIP
control
PP1H66K-PNUTSΔWBD
n=10222
I
Bru-seq DRB release 15 min.
control
PP1H66K-PNUTS
n=6817
J
Bru-seq DRB release 15 min.
20kb
control
PP1H66K-PNUTS
TOP1

Figure S4

**Supplemental Figure S4** Related to Figure 5

**PP1$^{H66K}$-PNUTS causes pol II CTD Ser5 hyperphosphorylation.**

**A.** Pol II Ser5-P ChIP seq metaplots as in Figure 5A (monoclonal 3E8) for a subset of genes with 2X enhanced antisense transcription as determined by Bru-Seq.

**B.** Pol II Ser5-P ChIP seq metaplots as in Figure 5A normalized to U snRNA signal.

**C, D.** Ser5-P ChIP-seq at the ACTG1 and RPL10A genes showing enhanced phosphorylation induced by PP1$^{H66K}$-PNUTS (blue arrows). Signal normalized to a mouse spike-in as in Figure 5C-E)

**E.** Ser5-P ChIP-seq at the snRNA gene RNU11 as in C, D showing little effect of PP1$^{H66K}$-PNUTS on Ser5 phosphorylation.

**F.** Metaplots of Ser5-P ChIP seq signal at histone genes in control (-dox) and PP1$^{H66K}$-PNUTS (+dox) expressing cells as in A. Note that in contrast to other protein coding genes PP1$^{H66K}$-PNUTS does not cause Ser5 hyperphosphorylation.

**G.** Metaplots of Ser5-P ChIP seq signal (monoclonal 4H8) in control (-dox) and PP1$^{H66K}$-PNUTS (+dox) expressing HEK293 cells showing hyperphosphorylation (blue arrow). Signals were normalized to U snRNA genes.

**H.** Metaplots of Ser5-P ChIP seq as in Figure 5G using a different monoclonal antibody (4H8).

**I, J.** PP1$^{H66K}$-PNUTS accelerates transcription. Bru-Seq 15 min after release from a DRB block in control (-dox) and PP1$^{H66K}$-PNUTS (+dox) expressing cells. Replicate of experiments in Figure 5H. Read counts in the two data sets were equalized by subsampling in H. Note the wave of transcription travels further with PP1$^{H66K}$-PNUTS (blue arrows).

### Figure S5

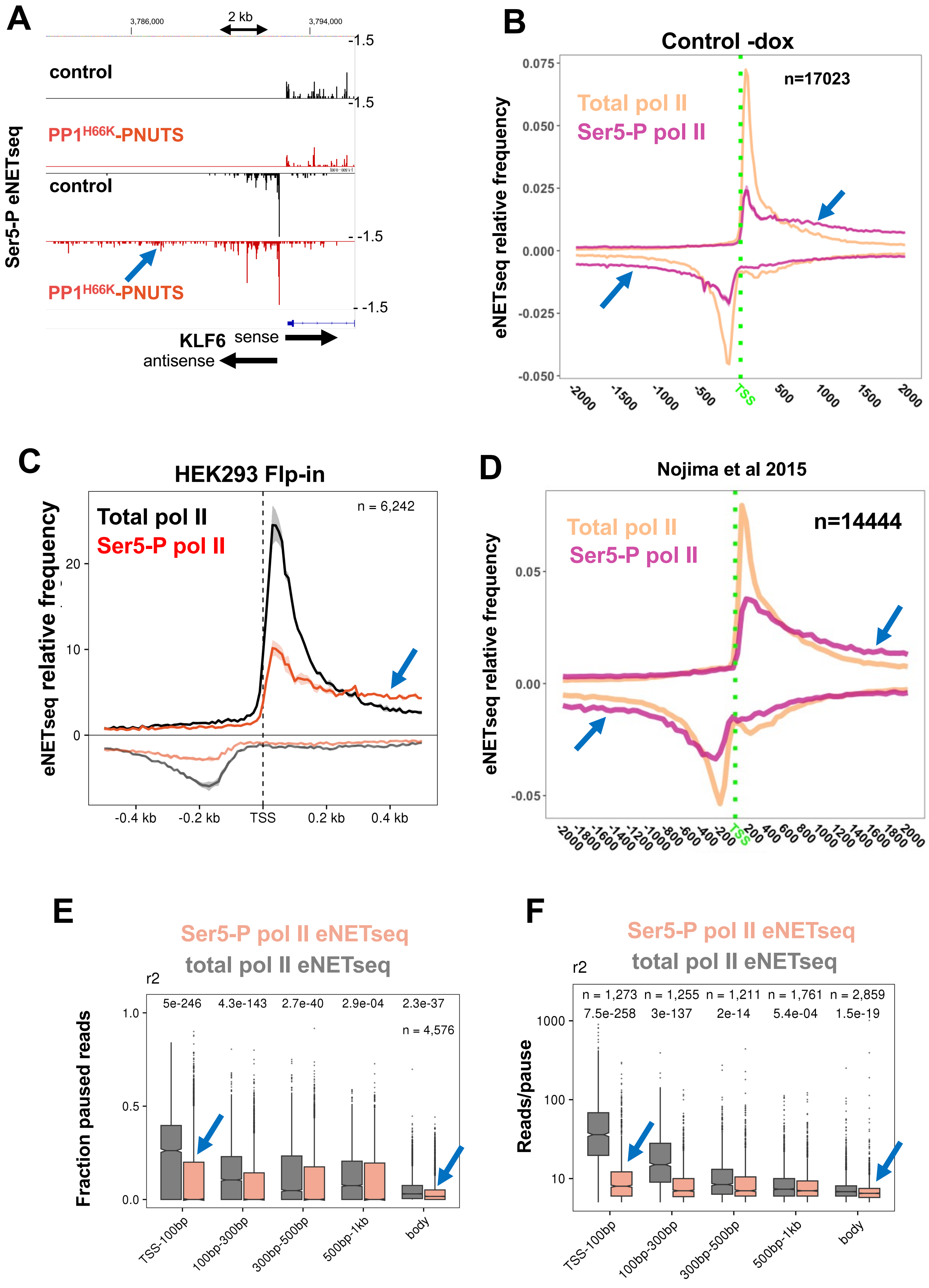

A
3,786,000
2 kb
3,794,000
1.5
control
-1.5
PP1H66K-PNUTS
control
-1.5
PP1H66K-PNUTS
-1.5
Ser5-P eNETseq
KLF6 sense
antisense
B
Control -dox
n=17023
Total pol II
Ser5-P pol II
eNETseq relative frequency
0.075
0.050
0.025
0.000
-0.025
-0.050
-2000
-1500
-1000
-500
TSS
500
1000
1500
2000
C
HEK293 Flp-in
Total pol II
Ser5-P pol II
n = 6,242
eNETseq relative frequency
20
10
0
-0.4 kb
-0.2 kb
TSS
0.2 kb
0.4 kb
D
Nojima et al 2015
Total pol II
Ser5-P pol II
n=14444
eNETseq relative frequency
0.05
0.00
-0.05
TSS
E
Ser5-P pol II eNETseq
total pol II eNETseq
r2
5e-246
4.3e-143
2.7e-40
2.9e-04
2.3e-37
n = 4,576
Fraction paused reads
1.0
0.5
0.0
TSS-100bp
100bp-300bp
300bp-500bp
500bp-1kb
body
F
Ser5-P pol II eNETseq
total pol II eNETseq
r2
n = 1,273
n = 1,255
n = 1,211
n = 1,761
n = 2,859
7.5e-258
3e-137
2e-14
5.4e-04
1.5e-19
Reads/pause
1000
100
10
TSS-100bp
100bp-300bp
300bp-500bp
500bp-1kb
body

**Supplemental Figure S5.** Related to Figure 6

**Enhanced elongation by Ser5-P pol II relative to total pol II.**

**A.** Ser5-P eNETseq at KLF6. Note elevated antisense signal (blue arrow) in cells expressing PP1$^{H66K}$-PNUTS (+dox) relative to control (-dox).

**B.** Metaplots of relative eNETseq signal as in Figure 6D (2 replicates) for Ser5-P pol II and total pol II in control PP1$^{H66K}$-PNUTS transduced cells (-dox). Note the relative amount of elongating versus 5' paused pol II is higher for Ser5-P pol II than total pol II (blue arrows).

**C.** Metaplots of relative eNETseq signal as in Figure 6D (2 replicates) for Ser5-P pol II and total pol II in HEK293 Flp-in cells. Note the relative amount of elongating versus 5' paused pol II is higher for Ser5-P pol II than total pol II (blue arrows).

**D.** Metaplots of relative mNETseq signal for Ser5-P pol II (GSE60358) and total pol II in Hela cells from ref. [47]. These data were generated using independent antibodies for total pol II and Ser5-P pol II than those we used. Note the relative amount of elongating versus 5' paused pol II is higher for Ser5-P pol II than total pol II (blue arrows).

**E. F.** Reduced pausing by Ser5-P pol II relative to total pol II in HEK293 Flp-in cells. Note reduced fraction of paused eNETseq reads and reads/pause for Ser5-P pol II both at the 5' end and within gene bodies (blue arrows). Pauses are positions with >5 reads where the signal is >3 SD above the mean in a 200 bp window. Reads were equalized by subsampling. (Replicates of the data shown in Figure 6E, F).
